# CLICK- chemoproteomics and molecular dynamics simulation reveals pregnenolone targets and their binding conformations in Th2 cells

**DOI:** 10.1101/2023.05.23.541892

**Authors:** Sougata Roy, Sudeep Roy, Bidesh Mahata, Jhuma Pramanik, Marco L. Hennrich, Anne-Claude Gavin, Sarah A. Teichmann

## Abstract

Pregnenolone (P5) is synthesized as the first bioactive steroid in the mitochondria from cholesterol. CD4+ and CD8+ immune cells synthesize P5 *de novo*, P5 in turn play important role in immune homeostasis and regulation. However, P5’s biochemical mode of action in immune cells is still emerging. We envisage that revealing the complete spectrum of P5-target proteins in immune cells would have multifold applications, not only in basic understanding of steroids biochemistry in immune cells but also in developing new therapeutic applications. We employed a CLICK-enabled probe to capture P5-binding proteins in live Th2 cells. Subsequently, using high-throughput quantitative proteomics we identified the P5-interactome in CD4+ Th2 cells. Our study revealed P5’s conserved mode of action in CD4+ and CD8+ immune cells. We identified novel proteins from mitochondrial and endoplasmic reticulum membranes to be the primary mediators of P5’s biochemistry in CD4+ and CD8+ immune cells. Applying advanced computational algorithms, we were able to generate near-native maps of P5-protein key molecular interactions that can lead to successful designing of novel molecular therapeutics strategies.

## Introduction

Pregnenolone (P5), the first bioactive steroid hormone of the steroid biosynthesis pathway, is synthesized in mitochondria from cholesterol. P5 is the progenitor of all glucocorticoids, mineralocorticoids, androgens, estrogens, and progesterone^1, 2^. Adrenal, gonads and placenta synthesize P5^3^. Besides, several steroidogenic cells have been reported to synthesize P5^4^. Role of P5 in the nervous system has been widely acknowledged^5, 6^. P5 enhances synapse and myelinization, induces growth of neurites^7^. P5 and its derivatives boost cognition and memory^6^ and also demonstrate therapeutic prospects in schizophrenia^8^. P5 in prostate cancer^9, 10^ and melanoma promotes tumour growth whereas in glioma it restricts tumour growth^11^. P5 mediates anti-inflammatory properties by activating degradation of proteins in the Toll-Like receptor signalling pathway^12^. Many cytoskeletal proteins are P5 receptors and regulate microtubule dynamics, cell migration and mitotic cell division^13, 14^.

The discovery of P5 as a lymphosteroid was reported recently by confirming its synthesis in lymphocytes and immune cells that infiltrate tumour^15, 16^. P5 synthesis in immune cells is a newly emerging domain that reveals the functional diversity of P5 across different cell types. In type 2 CD8+ cells local P5 synthesis drives its differentiation program^16^. CD4+ T cells are key mediators of immune responses against a plethora of infections^17^ and cancers^18^. Th2 lymphocytes are a primary subset of T cells that play key defensive roles to counter bacterial and helminth^19^ infections mounting adaptive immune responses^20, 21^. Th2 cells can synthesize P5, which in turn regulates their proliferation and class switching activity of B cells. In Th2 cells, P5 seems to play a crucial role in restoring immune homeostasis^15^.

Understanding P5’s regulatory action on Th2 immune cells becomes a topic of utmost importance. How does P5 regulate Th2 cellular biochemistry? To address this in our earlier study we used an interdisciplinary approach combining synthetic chemistry with high-throughput mass spectrometry. We designed and manufactured a synthetic and photoactivatable P5 analogue^22^. We captured P5 interactome in the cancer and CD8+ immune cells, the first proteome-wide P5-interactome map in any living cell. We identified 62 prospective target proteins of P5. The functional mapping of target proteins revealed a P5’s non-genomic mode of action. However, our analyses showed distinct pathways are targeted by P5 in cancer and CD8+ cells^22^. Th2 cell proliferation is inhibited by P5, however the underlying biochemical mechanism is missing.

Details of protein-ligand molecular interaction provides deep insight into the underlying mechanisms of bioregulation^23^. It also imparts novel structural information required for designing and development of novel drug molecules^24^. Molecular docking (MD) algorithms predict the mode and energy of binding in a ligand-target protein interaction. Molecular dynamics simulations not only improve MD prediction scores but also provide an atomic level resolution of the structure and dynamics of protein-ligand interactions. By combining these two tools near-native binding conformations can be achieved. P5-target protein binding mode, its structure and the underlying key interactions are unknown.

In this study, we have used the CLICK-enabled cell-permeable P5 probe^22^ in living Th2 cells. In conjunction with high throughput quantitative proteomics, we identified 11 ‘P5 binding’ target proteins in Th2 immune cells that are localized in the endoplasmic reticulum and mitochondria. Using an *in-silico* molecular simulation dynamics we present structural insight of the molecular interaction between P5 and its key targets. The P5 targets identified belong to some key pathways such as sterol biosynthesis, transport, protein processing and mitochondrial organization that clearly endorse the non-genomic role of P5’s in immune homeostasis. Most importantly, our results suggest that P5 activity in immune cells are mediated via mitochondria and ER. We envisage our study will not only provide a mechanistic understanding of the pregnenolone biochemistry in immune cells but also reveal the particulars of the binding efficiency and structural details of P5-target protein interaction.

## Results

### P5-C captures pregnenolone binding proteins from murine Th2 cells

P5 is vital in neurological^7, 25^ and cytoskeletal^13, 26^ milieu, however, its role in immune homeostasis independently or in the context of the tumour microenvironment is emerging (Figure 1a). We used the pregnenolone analogue P5-C, whose bioactivity, cell-permeability, and specificity as a mimic of native P5 has been described elsewhere^22^. To identify proteins enriched with P5-C in murine immune cells, we used the tandem mass tag (TMT) in conjunction with high throughput mass spectrometry for relative quantification. Figure 1b depicts the succinct plan of identifying P5-interactome from Th2 cells *in vivo*. To select for P5 specificity, P5-C capture was competed with 10X native P5. The proteins that are true P5-binding would be competed out when challenged with native P5. Only significantly enriched proteins that show effective competition when challenged with native P5 were selected. Subsequent analysis revealed 15 P5-interacting proteins enriched in Th2 cells (Figure 2a). All these 15 P5 binding proteins from murine Th2 (CD4+) are a subset of the 25 proteins identified to be P5-binding in the CD8^+^ T cells^22^ (Figure 2b). This reflects the functional conservation of P5 in CD8+ and CD4+ immune cells. To rule out any dual-binding proteins that might have been captured due to the protein’s affinity to the diazirine-alkyne linker, we took advantage of a recent study demonstrating the protein interacting with the diazirine-containing CLICK linker in mammalian cells^27^. While comparing, we found 4 prospective target proteins in our study that also showed dual binding affinity. Although it might be possible that these 4 proteins can bind P5 as well as diazirine, we decided to rule out these 4 common interactors thereby leading us to specific 11 proteins that are P5 targets in Th2 cells (Figure 2c).

**Figure 1.**
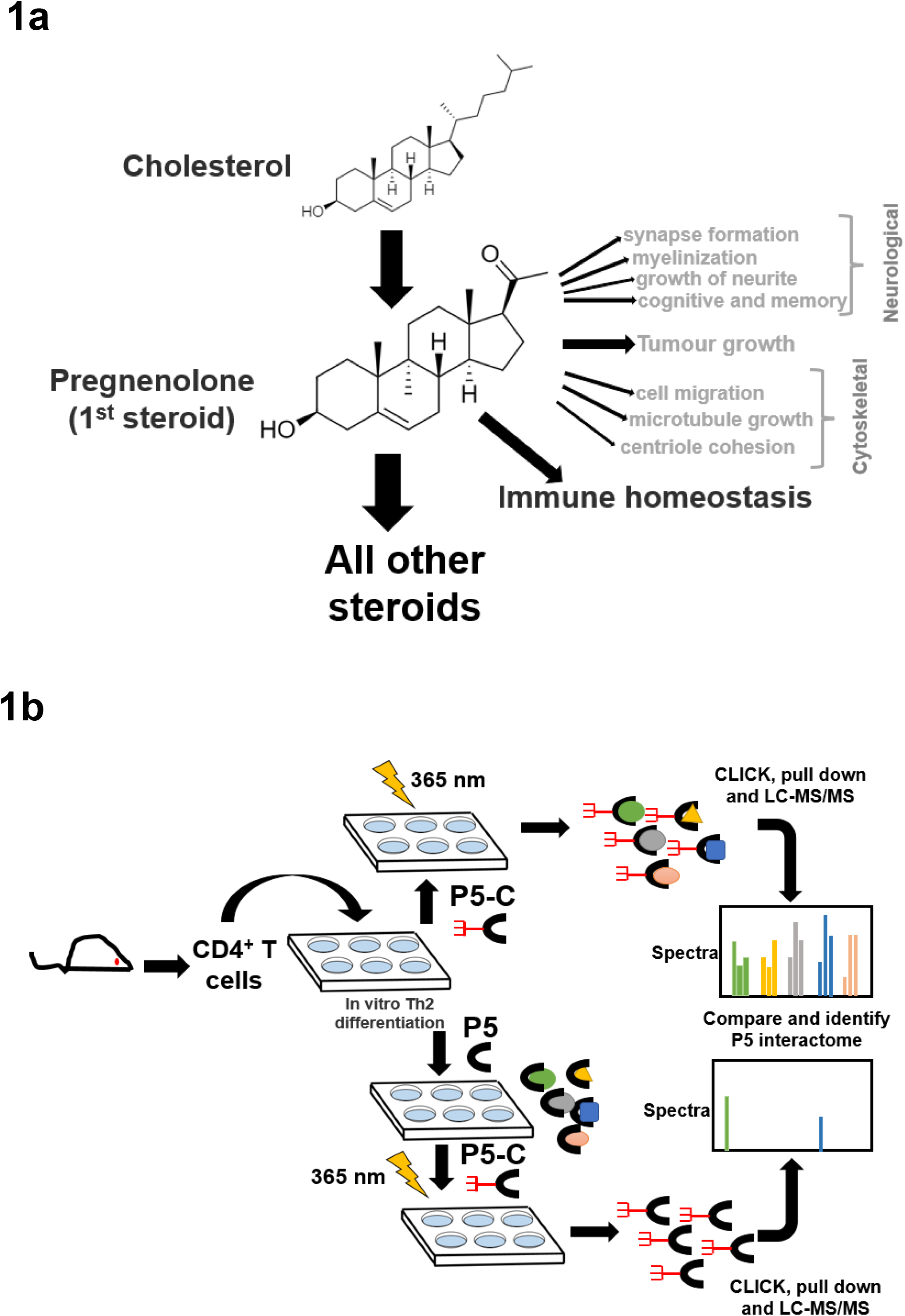
Pregnenolone (P5) and capturing its interactome in CD4+ Th2 cells. (A) The pleiotropic function of P5 is depicted in the above schematics. Its role in immune homeostasis has been highlighted that will be primary focus for our study here. (B) The schematics describes the capture of P5 interactome in live Th2 cells obtained from mice. The comparison of protein pulldown obtained with or without native P5 provides P5-specific binding proteins from live murine Th2 cells.

**Figure 2.**
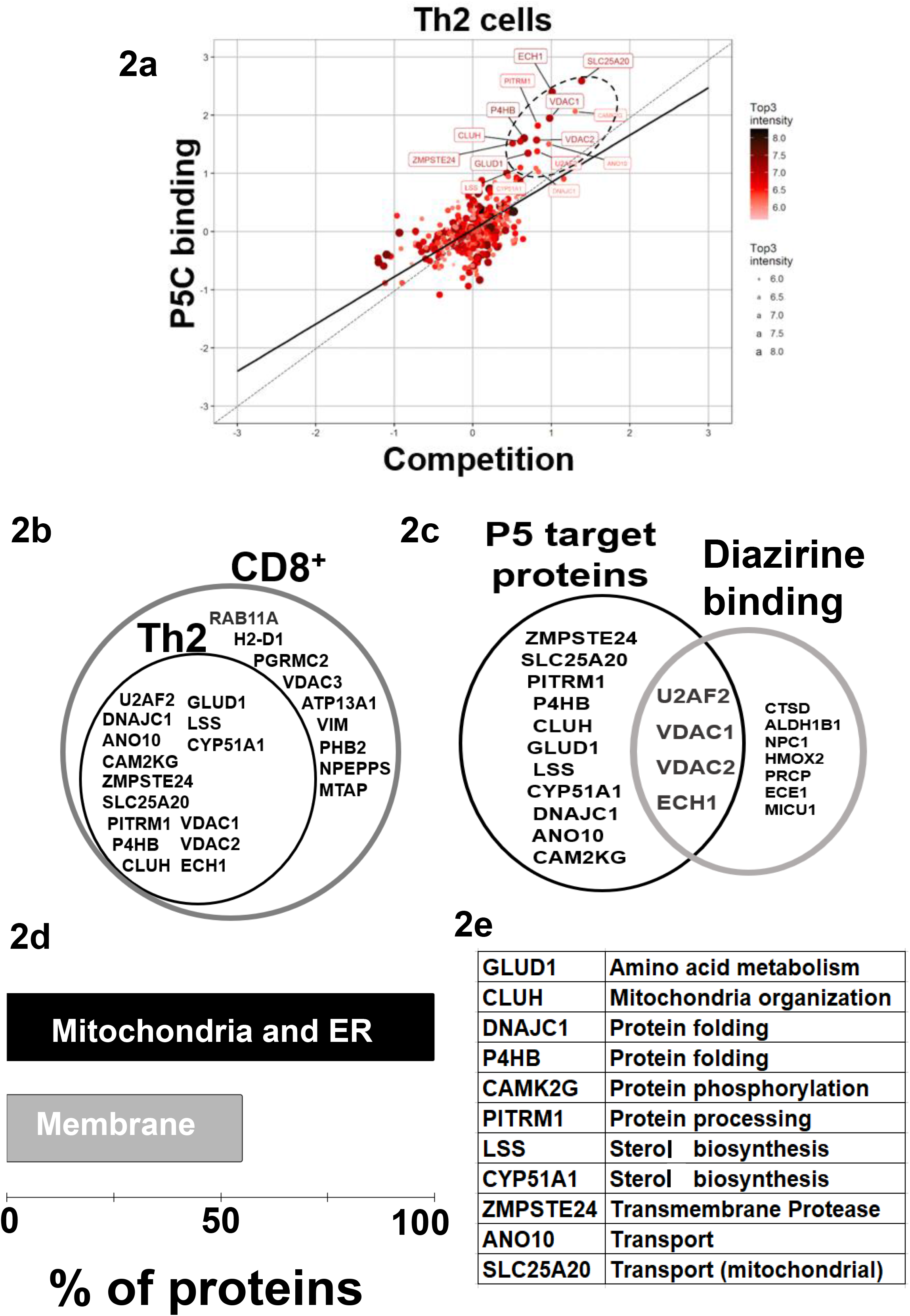
P5 interactome in Th2 cells and their functional analysis. (A) Fifteen proteins were significantly enriched as P5 binding in Th2 cells. The proteins extracted with P5-C either alone or after competition with P5. P5-C -interacting proteins were captured from live Th2 cells either in the presence of P5 (competition assay) or in its absence (experiment). The background (empty bead capture) were subtracted from the experiment and competition and then plotted on a log scale to identify P5 binding proteins. The solid straight line represent the regression equation fitting the two variables, and the dotted straight line demonstrate the X = Y linear relation. The solid circles in ellipse with dotted boundary contains the 15 P5-C binding proteins whose interaction can be competed out in the presence of native P5. (B) The diagrammatic representation of the overlap between the P5-binding proteins in two immune cell types, CD8+ T cells and CD4+ Th2 cells. All 15 P5-binding proteins captured in Th2 cells are a subset of proteins from CD8+ T cells. (C) The intersection region in the venn diagram shows four proteins that are known to bind both, P5 and diazirine. The 11 specific P5-binding proteins are encircled as “P5 target proteins”. (D) Functional categorization of the P5-binding proteins demonstrate a clear enrichment of ER and membranes (∼55%) that demonstrate the core functional pathway of P5 in immune cells. (E) The table enlisting the GO category ‘biological process’ of all the 11 P5-binding proteins in Th2 cells.

**Figure 3.**
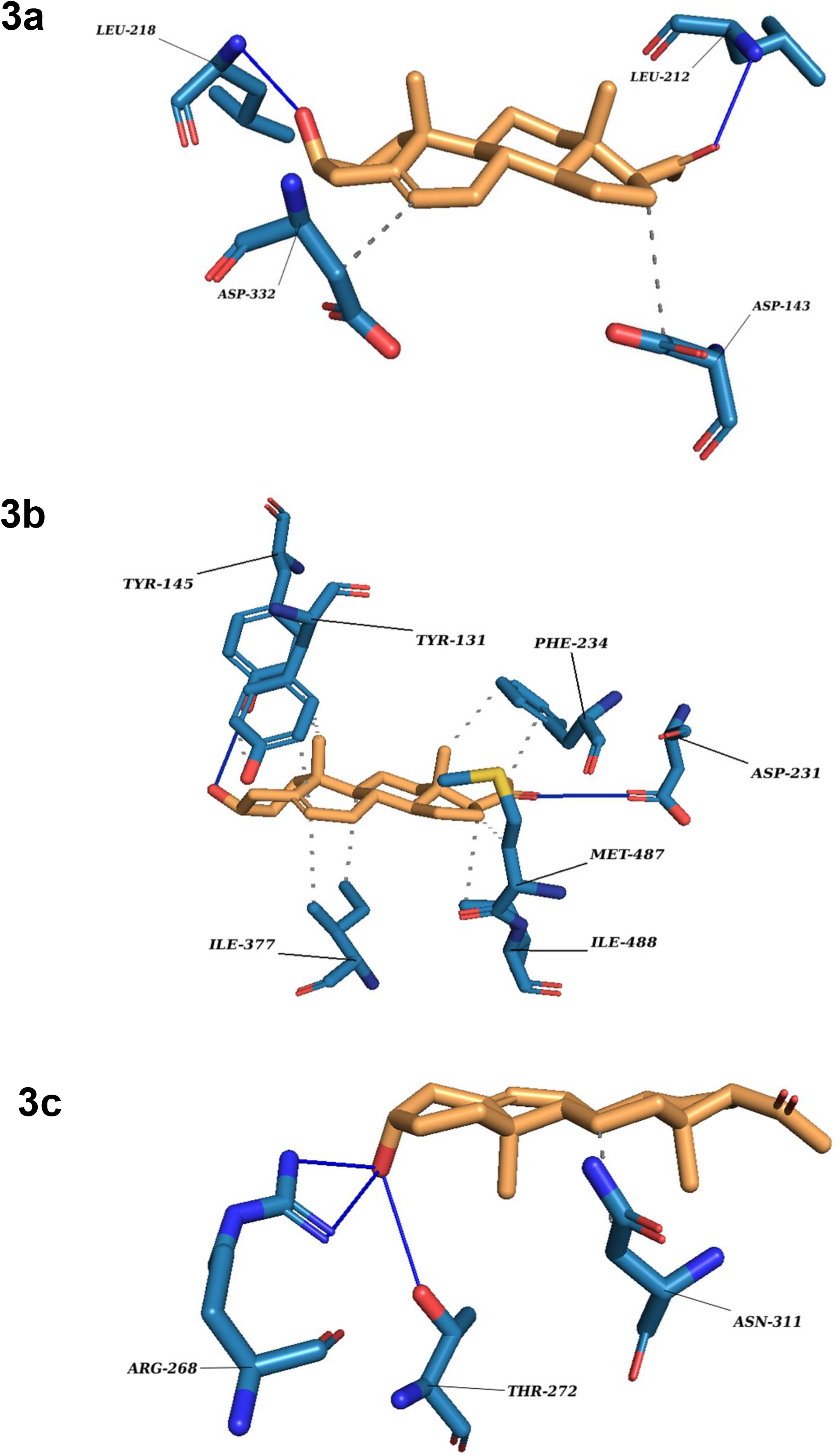
P5 docking complexes. (A) CLUH – P5 complex; Straight blue lines: hydrogen bonds; Dotted blue lines: hydrophobic interactions with the amino acids. (B) CYP51A1– P5 complex; Straight blue lines: hydrogen bonds; Dotted blue lines: hydrophobic interactions with the amino acids. (C) GLUD1– P5 complex; Straight blue lines: hydrogen bonds; Dotted blue lines: hydrophobic interactions with the amino acids.

**Figure 4.**
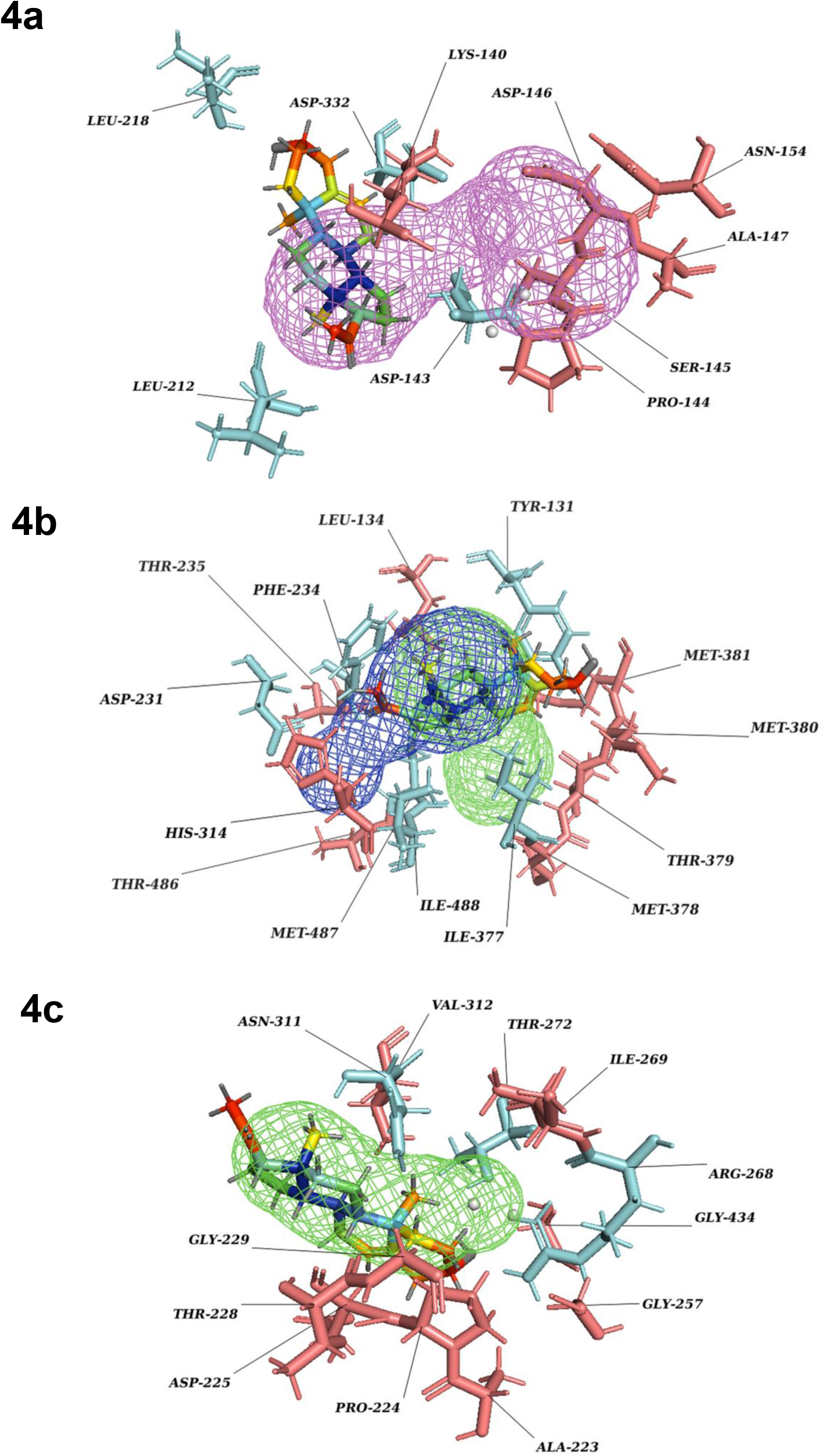
P5 transport through protein tunnel and channel. (A) Substrate P5 transport for CLUH protein through tunnel 6. Cyan color denotes hydrogen and hydrophobic interactions; Red Salmon color denotes other amino acids surrounding the tunnel. (B) Substrate P5 transport for CYP51A1 protein through tunnel 2. Cyan color denotes hydrogen and hydrophobic interactions; Red Salmon color denotes other amino acids surrounding the tunnel. (C) Substrate P5 transport for GLUD1 protein through tunnel 2. Cyan color denotes hydrogen and hydrophobic interactions; Red Salmon color denotes other amino acids surrounding the tunnel.

**Figure 5.**
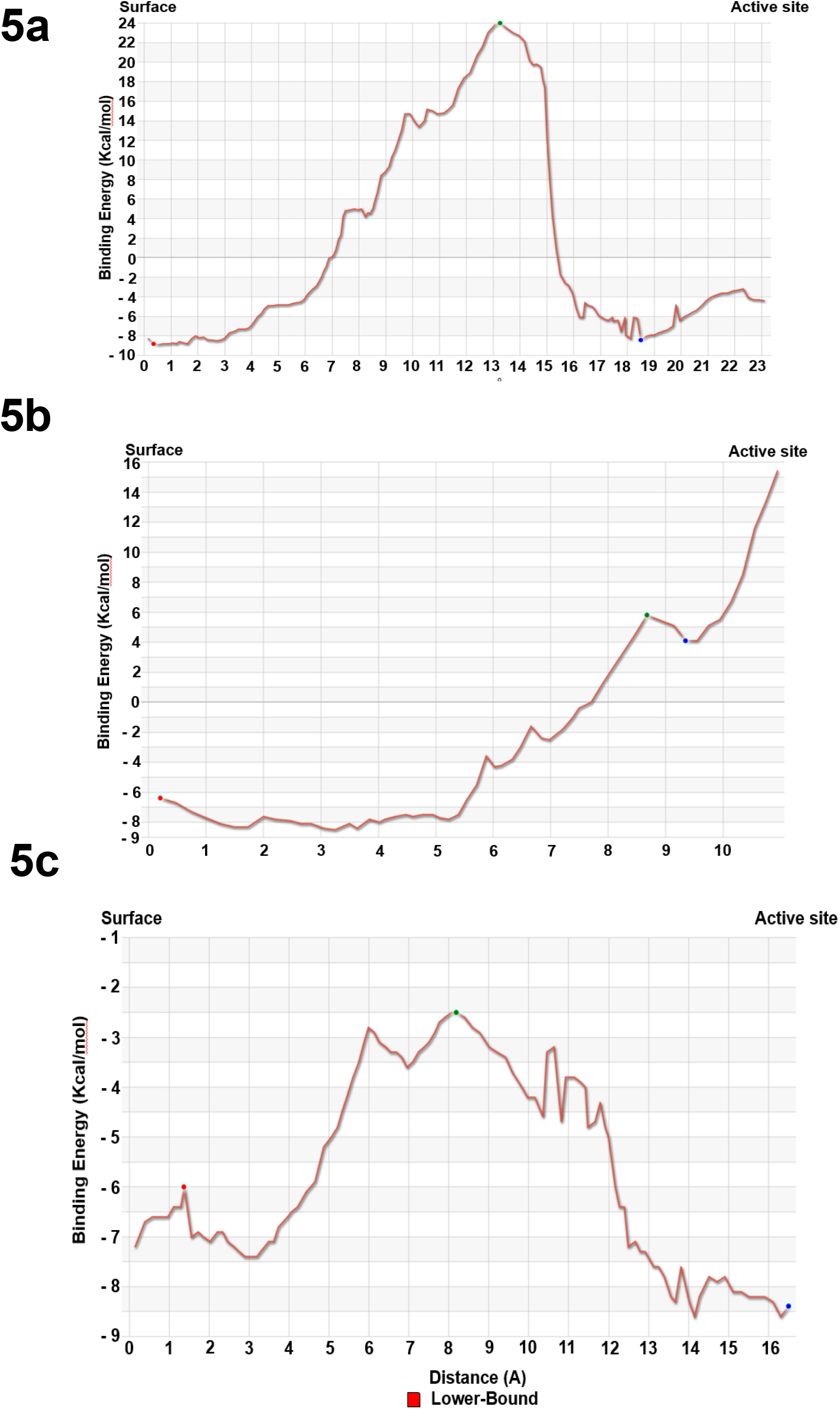
Energy profile of tunnel for proteins. (A) Energy profile of Tunnel 6 for CLUH protein [Red dot represents E Surface (kcal/mol), Green dot represents E Max (kcal/mol), Blue dot represents E Bound (kcal/mol)]. (B) Energy profile of Tunnel 2 for GLUD1 protein [Red dot represents E Surface (kcal/mol), Green dot represents E Max (kcal/mol), Blue dot represents E Bound (kcal/mol)]. (C) Energy profile of Tunnel 2 for CYP51A1 protein [Red dot represents E Surface (kcal/mol), Green dot represents E Max (kcal/mol), Blue dot represents E Bound (kcal/mol)].

### P5-Interacting proteins in Th2 cells are localised in mitochondria and endoplasmic reticulum

In agreement with our previous study^22^, P5-target proteins in Th2 are predominantly localised in ER, mitochondria, and the membranes. 9 of the 11 (81%) proteins are from the ER and/or mitochondria and 6 (55%) of these 11 proteins are on the membranes (Figure 2d). 2 proteins, PITRM1^28^ and SLC25A20^29^ are localized in the mitochondrion and GLUD1 is present in both ER and mitochondria respectively^30^. DNAJC1 and ZMPSTE24 are present in ER and nucleus. CLUH is the only cytoplasmic and ANO10 is the only plasma membrane protein in the P5-target list. The P5 interacting proteins are involved in cellular functions such as, mitochondrial organization, glutamate metabolism, protein folding, transport and steroid metabolism (Figure 2e).

### Molecular docking reveals how P5 interacts with its binding protein

To date, to the best of our knowledge structural features of P5 binding with its cognate binding partner proteins are obscure at molecular or sub-molecular level. We envisage it would be interesting to understand binding features and the conformational adaptation of the P5-target protein interaction. The substrate P5 is docked with CYP51A1, LSS, GLUD1, CLUH, P4HB and PITRM1 protein models through Induced Fit docking (IFD) to evaluate the binding mode. The most important feature of IFD is that the substrate, active site residues of the protein and its vicinity are considered as flexible for better assessment of protein-substrate interactions^31, 32^. The IFD results are provided in Table 1. The graphical representation in the form of protein-ligand interaction profiler is provided in Supplementary figure 1 (a-f) and detailed tabulated results in Table: ST1(a-f).

**Table 1:** Docking scores for selected protein models with P5

| Docking Scores |  | S_Score (kcal/mol) | Rmsd_refine (Å°) | E_conf | E_place | E_score1 | E_refine | E_score2 |
| --- | --- | --- | --- | --- | --- | --- | --- | --- |
| P5 |  |  |  |  |  |  |  |  |
|  | CLUH | -7.0705 | 1.2121 | 20.3313 | -41.7799 | -7.4497 | -38.7172 | -7.0705 |
|  | CYP51A1 | -6.9349 | 1.4454 | 16.3240 | -60.3607 | -9.0636 | -31.0231 | -6.9349 |
|  | GLUD1 | -6.9486 | 1.6890 | 18.6725 | -82.1230 | -9.7515 | -29.2101 | -6.9486 |
|  | LSS | -6.0083 | 1.5237 | 14.9390 | -69.0326 | -9.1873 | -27.3442 | -6.0083 |
|  | P4HB | -6.2503 | 1.1224 | 44.9485 | -59.6411 | -9.4770 | -19.3375 | -6.2503 |
|  | PITRM1 | -6.0898 | 1.6296 | 18.0037 | -67.0599 | -8.3369 | -27.9744 | -6.0898 |

The results are obtained under the following criteria. **S** score is the score of the final stage of the docking run. **RMSD** defines the deviation between the pose, in Å, from the original ligand. **RMSD refine** defines the deviation between the poses before refinement and the poses after the refinement stage. **E_conf** as the energy of the conformer at the end of the refinement. Energy is calculated with the solvation option set to Born. **E_place** as the score from the placement stage. **E_score1** and **E_score2** are the scores for Rescoring stages 1 and 2 respectively. **E_refine** is the score from the refinement stage. It is calculated to be the sum of the vander Waals electrostatics and solvation energies, under Generalized Born solvation model (GB/VI). Lower final S-scores indicate more favourable poses. The detailed docking result is provided in Table: 1.

The docking scores range from -7.0705 to -6.0083 (kcal/mol) for all the set of protein-substrate complexes. The RMSD values are within the permissible limits. The docking scores obtained are comparable and the substrate binds with the proteins. CLUH, CYP51A1 and GLUD1 show better binding affinity according to the docking results. The findings are further validated through molecular simulations and pose rescoring with MM-GBSA calculations.

### Evaluation of stability of protein models and their thermodynamic properties

The docked protein-substrate complex was further subjected to molecular dynamic simulations to validate the docking results. Various thermodynamic properties and stability parameters were evaluated for protein-substrate complexes. The Desmond module from Schrodinger’s Maestro Suite was implemented for further validation. Several thermodynamic properties for protein and substrate, such as Root mean square deviation (RMSD), Root mean square fluctuation (RMSF), Radius of Gyration (ROG), Secondary structure element (SSE), Protein-Substrate timeline and Protein-Substrate contacts of CYP51A1, LSS, GLUD1, CLUH, P4HB and PITRM1 substrate complexes were monitored for 100 ns MD simulation run.

#### A) Protein RMSD

Protein RMSD values should be within the range of 1-4 Å which indicates that the simulation has equilibrated. Its fluctuation towards the end of the simulation is around thermal average structure. The RMSD values recorded is quite high for all the proteins. We expect higher RMSD values of homology derived model proteins. The order of protein RMSD values are CYP51A1 (8.8 Å)>CLUH (9.0 Å)>P4HB (12.8 Å)>GLUD1 (13.5 Å)> PITRM1 (21.0 Å)> LSS (27.0 Å). (Fig S1 (a-f)).

#### B) Substrate RMSD

The value of substrate RMSD is less than protein RMSD and has stabilized at the end of the simulation for CLUH, CYP51A1, GLUD1 and P4HB proteins-substrate complex. This indicates that the substrate is stable with respect to the protein and its binding pocket. If the RMSD values for the substrate observed are significantly larger than the RMSD of the protein, then it is likely that the substrate has diffused away from its initial binding site. This exception is observed in PITRM1 where the RMSD value of substrate is slightly higher than protein backbone. The order of substrate RMSD values are CLUH (4.1 Å)>CYP51A1 (4.4 Å)>GLUD1 (5.0 Å)>P4HB (4.0 Å) >LSS (19.0 Å)>PITRM (22.0 Å). The detail figure is provided in Fig S1 (a-f).

#### C) Protein RMSF

The Root Mean Square Fluctuation (RMSF) is useful in characterising local changes along the protein chain. On the plots, peaks indicate areas of the protein that fluctuate the most during the simulation. It is observed that the N-terminal fluctuates more than the C-terminal and other parts of the protein. Secondary structural elements like alpha helices and beta strands are usually more rigid than the unstructured part of the protein, and thus fluctuate less than the loop regions. Our finding suggests that almost all the proteins have high values of fluctuation in the N- and C-terminals. The fluctuation was slightly less in CYP51A1 and LSS for C-terminal due to comparative stable Super Secondary Elements (SSEs) of the proteins (Fig S2 (a-f)).

##### Substrate Contacts

Protein residues that interact with the substrate are marked with green-coloured vertical bars.

#### D) Substrate RMSF

Substrate RMSF shows the substrate’s fluctuations broken down by atom, corresponding to the 2D structure. The substrate RMSF may give you insights on how substrate fragments interact with the protein residues and their entropic role in the binding event. Under the top panel, the **’Fit on Protein’** line shows the substrate fluctuations, with respect to the protein. Our finding shows that LSS and PITRM1 have high entropy values. We assume this is due to formation of several weak hydrogen bonds and hydrophobic interactions of LSS and PITRM1 proteins with the substrate. The protein-substrate complex is first aligned on the protein backbone and then the substrate RMSF is measured on the substrate heavy atoms (Fig S3 (a-f)).

#### E) Secondary structure elements

Protein secondary structure elements (SSE) like alpha-helices and beta-strands are monitored throughout the simulation run. The plot provided in Fig S4 (a-f), summarises the SSE composition for each trajectory frame over the course of the simulation, and the plot at the bottom monitors each residue and its SSE assignment over time. Our finding suggests that protein structure had average confirmation with 49.03% (CLUH), 55.01% (CYP51A1), 49.52% (GLUD1), 45.13% (LSS), 46.54% (P4HB), 52.96% (PITRM1) SSE, mainly composed of helices and strands rather than loops and turns that showed conformational changes during MD simulations run (Fig S4 (a-f)).

#### F) Radius of Gyration

ROG is analysed to examine the compactness of the model protein. All the proteins were slightly destabilized throughout the run and never stabilizes to attain compactness till the end of the simulation. (Fig S5 (a-f)).

#### G) Protein-Substrate Contacts

Protein interactions with the substrate can be monitored throughout the simulation. These interactions can be categorized by type and summarized, as shown in (Fig S6 (a-f)). Protein-substrate interactions (or ’contacts’) are categorized into four types: Hydrogen Bonds, Hydrophobic, Ionic and Water Bridges. The stacked bar charts are normalized over the course of the trajectory: for example, a value of 0.7 suggests that 70% of the simulation time the specific interaction is maintained. Values over 1.0 are possible as some protein residue may make multiple contacts of the same subtype with the substrate.

Our finding suggests that LEU212 (strong), LYS216 (weak), LEU218 (strong), LYS303 (weak) and ASP332 (weak) interacts with pregnenolone in form of H-bonds for CLUH protein. HIS314 (weak), ARG382 (very strong) and HIS447 (strong) forms H-Bonds with pregnenolone for CYP51A1 protein. Similarly, TYR131 and PHE234 forms strong hydrophobic interactions for CYP51A1. LYS147 (strong), GLY148 (weak), THR256 (weak), ARG268 (weak), GLY434 (weak) and SER438 (partially strong) forms H-Bonds with pregnenolone for GLUD1 proteins. In case of LSS, ASP148 (weak), ASN279 (weak), TYR287 (partially strong) and ARG640 (weak) forms overall weak hydrogen bonds. Similarly, for PITRM1, ASN136 (weak), TYR143 (weak), TYR380 (weak), GLY382 (weak), TYR383 (weak), THR446 (weak) and SER453(weak) forms overall weak H-bonds. P4HB forms the strongest H-bond amongst all the proteins reported with the participation of LYS232 (weak), GLN235(weak), LEU238 (strong), GLY286 (weak), ILE288(weak) and PHE290 (strong). It is to be noted that LEU236 and LYS285 participates in strong hydrophobic interactions for P4HB with the substrate. Several other bonds and interactions such as Ionic bond, hydrophobic interactions and water channels are also reported for all the proteins-substrate complexes under investigation. The results were concluded on the basis of protein-substrate contacts bar diagram and residue decomposition values. There were small instances where residue decomposition analysis fails to detect weak hydrogen bonds for LSS and PITRM1 proteins in pose-rescoring studies. We have considered the bonds keeping in mind all the possible poses generated during the simulation run. The detailed representation is provided in (Fig S6 (a-f)), Table: ST2 and Residue Decomposition Complete.xls.

#### H) Protein-Substrate timeline

A timeline representation of the interactions and contacts (H-bonds, Hydrophobic, Ionic, Water bridges) is summarised (Fig S7 (a-f)). The top panel shows the total number of specific contacts the protein makes with the pregnenolone over the course of the trajectory. The bottom panel shows which residues interact with the substrate in each trajectory frame. Some residues make more than one specific contact with the pregnenolone, which is represented by a darker shade of orange, according to the scale to the right of the plot (Fig S7 (a-f)).

##### Pose rescoring with MMGBSA

The confirmations of the docking results and substrate efficiency were determined by calculating binding free energy through Prime-MM/GBSA (Molecular Mechanics/Generalized Born Surface Area) (Prime, Schrodinger, LLC, New York, NY. 2022). MM-GBSA scoring usually provides a significant correlation with experimentally determined data. The calculated binding free energy for the docked complexes using the OPLS-AA force field and the GBSA continuum solvent are represented in Table 2. The binding free energy for the protein-substrate range from -27.29kcal/mol to -58.35kcal/mol. The substrate molecule is considered as the best that utilises less energy to comfort inside the active pocket of the protein. In our study CLUH, CYP51A1 and P4HB showed better rescoring binding energy values. The results are an improvement over induced fit docking results where the finding shows that CLUH, CYP51A1 and GLUD1 showed better binding affinity values. MM/GBSA calculation classified the binding affinity of the substrate in the exact place as docking energy while there were little changes in the order for intermediate docking energy holding compounds (Table 2).

**Table 2:** MM-GBSA values for protein models with P5

| S. No. | Proteins | $\Delta G$ Bind | $\Delta G$ Bind Coulomb | $\Delta G$ Bind vdw | $\Delta G$ Bind Covalent | $\Delta G$ Bind Solv GB | $\Delta G$ Bind Lipo | $\Delta G$ Bind H Bond |
| --- | --- | --- | --- | --- | --- | --- | --- | --- |
| 1 | CLUH | -58.35616818 | -12.22089589 | -47.73719881 | 0.452964344 | 18.88040592 | -16.85542031 | -0.876023442 |
| 2 | CYP51A1 | -57.6212031 | -8.961990524 | -48.5034085 | 1.316226552 | 25.35348989 | -26.06089958 | -0.764620935 |
| 3 | GLUD1 | -29.17202635 | -4.748506696 | -36.82294515 | 1.397948969 | 22.92806308 | -11.28422616 | -0.642360396 |
| 4 | LSS | -28.56342559 | -4.33566621 | -24.56548335 | 0.760308397 | 11.32898609 | -11.53867904 | -0.212891473 |
| 5 | P4H8 | -57.04931056 | -17.12902166 | -46.02070591 | 1.835749835 | 23.05314964 | -17.71682611 | -1.071656346 |
| 6 | PITRM1 | -27.29123782 | -4.965643417 | -2540774676 | 0.66446038 | 15.3832475 | -12.73605131 | -0.229504201 |

**Table3:** Energy profile of substrate transport within the tunnel for GLUD1, CYP51A1 and CLUH protein-substrate complex

| Substrate | Tunnel | Direction | $E_{\text{BOUND}}$<br>[kcal/mol] | $E_{\text{MAX}}$<br>[kcal/mol] | $E_{\text{SURFACE}}$<br>[kcal/mol] | $E_a$<br>[kcal/mol] | $\Delta E_{\text{BS}}$<br>[kcal/mol] |
| --- | --- | --- | --- | --- | --- | --- | --- |
| GLUD1_P5 | 2 | IN | 4.1 | 5.8 | -6.4 | 12.2 | 10.5 |
| CYP51A1_P5 | 1 | IN | -7.3 | 3.5 | 1.5 | 2.0 | -8.8 |
| CYP51A1_P5 | 2 | IN | -8.4 | -2.5 | -6.0 | 3.5 | -2.4 |
| CLUH_P5 | 6 | IN | -8.4 | 24.0 | -8.8 | 32.8 | 0.4 |

Finally, to measure the contribution of the individual residues, the MM/GBSA decomposition analysis was performed. MD simulation analyses of the substrate−residue interactions in the above section are based on the trajectory averaged and minimised, i.e., on static protein-substrate conformations, the MD-based decomposition considers the contribution of protein residues in all possible binding modes. The detailed result is provided in an excel sheet provided in the Supplementary data (Residue Decomposition Complete.xls).

CaverDock is an in-house tool that uses Caver, to identify tunnels in protein structures, and an optimised version of the well-established algorithm from AutoDock Vina to calculate possible ligand trajectories along those tunnels and the corresponding binding energies. CaverDock discretises each identified tunnel into a series of discs and models a ligand’s passage through the tunnel by constraining one ligand atom to lie within a disc, sequentially. The ligand’s conformation and binding energies are then calculated using Autodock Vina, with the ligand (aside from the constrained atom) being free to explore the conformational space; the protein is treated as a rigid body. Once the conformation and binding energy have been calculated, the constrained atom is shifted to the next disc and the process is repeated until the ligand has moved through the full length of the tunnel. The tool is previously successfully utilized in various studies, including spike glycoprotein of SARS-CoV2^33^, and leukotriene^34^. Our Caver study confirms that substrate pregnenolone interacts well with important bottleneck amino acid residues in the tunnel with GLUD1, CYP51A1 and CLUH proteins (Fig: 2(a-c) and Table: ST3). Bottleneck residues are marked (salmon-red) in bold for the respective proteins are involved in the interaction with pregnenolone. Tunnel 2(GLUD1-ARG268, THR272, ASN311), Tunnel2 (CYP51A1-TYR131, ASP231, PHE234, ILE377 and ILE488) and Tunnel 6 (CLUH-ASP143) shows hydrogen bonds and hydrophobic interactions with the pregnenolone (Table: ST4). The energy profile of the tunnels for respective enzymes have been performed by selecting the correct tunnel that allows substrate to transport. We can observe that after passing through the bottleneck residues, pregnenolone shows selective binding at the active site for CYP51A1 and CLUH enzyme with a favourable binding energy of -8.4 kcal/mol for both the proteins respectively. The energy barrier around the gateway residues were lower for both CYP51A1(-8.8(kcal/mol-Tunnel 1) and (-2.4(kcal/mol-Tunnel 2) and CLUH (0.4 kcal/mol-Tunnel 6) respectively (Table: 3). Pregnenolone interaction with GLUD1 show slightly higher binding energies (4.1 kcal/mol). The difference of the binding energy of pregnenolone in the active site and at the surface for the gateway residues for Pregnenolone-GLUD1 complex was recorded high (10.5 kcal/mol). The detailed CaverDock result in form of energetic profile are provided in Fig: 3(a-d). The substrate doesn’t pass through tunnels for proteins LSS, P4HB and PITRM1 (Fig: S8(a-c)) and the binding of pregnenolone with the amino acids takes place outside the tunnels.

## Discussion

Although P5’s importance as a bioactive molecule with pleiotropic functions is beyond doubt, its biochemical mode of action is not fully understood. P5 demonstrates crucial functions in regulation of inflammation and immunity^12, 15, 16, 35^. CD4+ Th2 and CD8+ subtypes of immune cells produce P5 via local steroidogenesis^15, 16^. We have deciphered P5 - interactome in the CD8+cells^22^, P5-interactome in Th2, a CD4+ subtype would reveal conserved and specific pathways of P5 biochemistry in these immune cells. With this aim we sought out to find P5-interacting proteins specific to Th2 cells. We envision that our finding could provide a comprehensive biochemical underpinning of P5 that are concurrent and distinct to immune cell-types.

The bioactive and cell permeable CLICK-based P5 analogue^22^ was used to mimic native P5 in live Th2 cells. In conjunction with high throughput mass spectrometry, we revealed a comprehensive proteome-wide interactome-map of P5 in Th2 immune cells. We identified 11 P5 binding proteins in Th2 cells. as specific P5-interacting proteins Interestingly, all these 11 proteins are a subset of the 25 P5-binding proteins that were previously identified in CD8+ cells by us^22^. This suggests conservation of key P5 function across CD4+ and CD8+ immune cell types. We envisage that these proteins may represent the core conserved functional pathways regulated by P5 in immune cells. The identified proteins were from mitochondria and endoplasmic reticulum (>80%) and membranes (∼55%) (Figure 2d), which demonstrated a non-genomic mode of P5 activity corroborated by many previous studies^5, 13, 14, 22, 26^.

Mitochondria^36^ and ER^37^ are known to play crucial roles in immune cells. P5 target proteins localized in these organelles suggests P5’s regulatory role in modulating Th2 cells. In our previous study we found mitochondria acyl-carnitine transport pathway proteins as P5-targets in CD8+ cells^22^. In CD4+ cells we found the mitochondrial SLC25A20, the transporter of the acyl-carnitine, to be a target of P5. P5 is synthesized in mitochondria^2^, we found clustered mitochondria protein homolog (CLUH) to be a target of P5. CLUH is an RNA-binding protein that controls mitochondrial biogenesis and distribution by modulating the expression of nuclear-encoded mitochondrial RNAs^38^. It is therefore possible that P5 can regulate mitochondrial homeostasis via CLUH. Understanding the structural details of P5-CLUH molecular interaction can provide useful insight of this function. Mitochondrial Glutamate dehydrogenase 1 (GLUD1), another P5 target, is known to play key role in replenishing TCA cycle intermediate ɑ-ketoglutarate from L-glutamate^39^. ɑ-ketoglutarate performs crucial role in T cell differentiation^40^. Furthermore, GLUD1 enzyme plays an important role in insulin balance^41^ and is one of the top candidates that is involved in long term memory and spatial learning^42^. P5’s role in the nervous system is well-known^5^ and GLUD1 may be the entry point to comprehend the biochemical pathway of P5 activity in brain cells. Steroid and their derivatives have a variety of roles in regulation of immunity mediated by the different immune cell types^43^. Their roles are still emerging, in this context, we found two key enzymes CYP51A1, and LSS in the sterol biosynthetic pathway to be P5 targets. This suggests a regulatory role of P5 in the biosynthesis of sterol, including cholesterol, the immediate parent molecule from which P5 is synthesized. P5 regulating its own homeostasis within steroidogenic cells via feedback inhibition seems plausible. Other targets of P5 are related to cellular and sub-cellular transport and protein processing and folding. All these functions are crucial in regulation of the immune system.

We further provided insight into the molecular interaction of P5 and its key targets at the atomic level. To do this we employed state-of-the art docking, molecular dynamics, tunnel and channel detection tools. The docking results clearly indicate binding up of proteins with the substrate pregnenolone. The values are comparable and the RMSD values are within the range of 2Å. CLUH, CYP51A1 and GLUD1 protein shows better binding affinity with the substrate pregnenolone. The docking results are well complemented by the simulation findings. RMSD values of the modelled protein backbone are higher but the substrate fits the binding site. The NPT ensemble system has equilibrated till the end of the simulation run. The exception was recorded for PITRM1 protein (Fig S1(a-f)). Protein RMSF was recorded for all the proteins. The property is useful in characterizing the local changes along the protein chain. All the proteins under investigation constitute mainly of alpha helices and beta sheets. CYP51A1 and LSS recorded the least fluctuation of amino acids amongst all the proteins under investigation (Fig S2(a-f). The substrate RMSF values for proteins LSS and PITRM1 have high entropy values. This is due to the formation of several weak hydrogen bonds and hydrophobic interactions (Fig S3(a-f)). All the proteins did not attain the compactness throughout the simulation run (Fig: S5(a-f)). Substrate-protein contacts clearly show that P4HB forms the strongest H-bonds amongst all the protein-complex under investigation. It is followed by CYP51A1 and CLUH proteins respectively (Fig S6 (a-f)) and Table: ST2. MM-GBSA studies were performed to study the correlation of *in-silico* findings with experimentally determined data. The simulation analysis performed is based on the trajectory averaged and minimised, i.e., on static protein-substrate conformations. MM-GBSA considers the trajectory output generated in the simulation run to further validate the docking findings. The main purpose of the study is to look deep into the energy values that utilizes less energy for the substrate to comfort inside the active pocket of the proteins. The study reveals that CLUH, CYP51A1 and GLUD1 showed better binding affinity values while taking into special consideration of intermediate docking energies (Table 2). Finally, we measured the contribution of the individual residues in the interaction with substrate through Residue Decomposition Analysis. This analysis considers the contribution of protein residues in all possible binding modes. The result provides deep insight into the possible bond formation of the substrate complexes in the form of Binding energies, Coulombic and hydrogen bonds, Van der Waals, Lipophilic & hydrophobic interactions (Residue Decomposition Complete.xls).

Protein forms functionally important local substructures, such as active sites, allosteric sites, tunnels and channels. The tunnels are mainly present in globular proteins with catalytic function (enzymes) and serve as the access pathways for substrates, products, co-factors, water molecules and /or inhibitors from a bulk solvent to buried active sites. Our study reveals that pregnenolone interacts with important bottleneck amino acid residues in the tunnel with GLUD1, CYP51A1 and CLUH proteins. The energy profile analysis for the tunnels for respective proteins have been performed to select the correct tunnel that allows substrate to transport (Table: 3). The various parameters are summarized as: **Tunnel** – selected protein tunnel; **Substrate** – name of the used substrate; **Direction** – direction of the CaverDock calculation; **E_Bound_** – the binding energy of the substrate located in the binding site; **E_Max_** – the highest binding energy in the trajectory; **E_Surface_** – the binding energy of the substrate located at the protein surface; **E_a_** – activation energy of association, **E_Max_ - E_Bound_** for products, **E_Max_ - E_Surface_** for reactants (describes the difficulty of getting through the tunnel; kinetics); **ΔE_BS_** – difference of the binding energies of the substrate in the active site and at the surface (corresponds to enthalpy; thermodynamics).The CaverDock result shows the energetic profile of the substrate binding. The graph generated for the respective tunnel allows us to select the values that correspond to the bound state (E_bound_), maximum energy (E_max_) and unbound state (E_surface_). After selecting these three points CaverDock automatically calculates the estimate of the binding energy barrier (E_A_) and the difference in energy of the bound and unbound states (E_BS_). To study the ease of access to the druggable site for the substrate, E_max_ (the highest binding energy in the trajectory) and Ea (activation energy of association: E_Max_-E_Surface_ of the reactants) are two parameters of importance. To select well binding substrates, we should aim for the lowest values in both E_max_ and E_a_ in terms of the energy values. In our study, for GLUD1 and CLUH, only one tunnel is being reported for substrate transport. In case of CYP51A1, two tunnels are reported. It can be seen from Table:3 that the values of E_a_ are comparable for both the tunnels but Tunnel 2 is comparatively stable for CYP51A1 to transport substrate than Tunnel 1 due to lower value of E_max_. The detailed representation of the energy profile for the selected protein-complex is provided in Fig: 3(a-d). The substrate doesn’t pass through tunnels for LSS, P4HB and PITRM1 protein-complexes (Fig: S8(a-c)). The binding of substrates for the above-mentioned proteins takes place outside the active sites.

P5’s role as a lymphosteroid has not been investigated in detail. Our study not only revealed the specific P5-interacting proteins in live murine Th2 cells but also identified near-native key P5-protein molecular interactions. We found mitochondrial and ER localised proteins play key roles in mediating P5 activity in CD4+ and CD8+ immune cells. Some of the P5-binding proteins identified from our study, such as GLUD1, PITRM1, P4HB and CYP51A1 are already well-known therapeutic targets for drug development. We believe our study has unlocked a new domain to explore the mechanism of P5’s mode of action in immune cells that will provide insights into designing innovative drug targets from the molecular dynamics output of P5-protein interactions.

## Materials and Methods

### Materials

The P5 analogue P5-C has been described previously. The probe is bioactive, cell permeable and has the capacity to specifically capture P5 binding proteins.

### Th2 Cell Proliferation

The detailed method is described previously^22^.

### P5 Binding proteomics on murine Th2 cells

10 million murine CD4+ Th2 immune cells were used in three different experiments with two replicates for each. In the first experiment, Th2 cells in phenol free RPMI were incubated with only the vehicle (without probe), second experiment had 10 µM P5-C and the third was incubated with 100 µM (10X) of P5 before 10 µM of P5-C incubation. The following protocol was the same for all the experiment and its replicates. The cells were centrifuged, and the media was removed, cells were washed once with cold PBS and then irradiated at 365nm for 15 minutes in 200 μL of cold PBS at 4°C. Subsequently, the PBS was removed after centrifugation and the cells were washed once with ice cold PBS. Lysis of Th2 cells were done in PBS containing 0.2% SDS. Lysates were centrifuged and the supernatant containing the clear lysates were used for copper-click and affinity purification. Following 3 times PBS/0.1% SDS, the beads were washed 3 times with 1 mL of PBS. The washed neutravidin beads were sent to EMBL proteomics core facility for TMT labelling, peptide fractionation and mass spectrometry.

### Mass Spectrometry

#### Sample preparation and TMT labelling

The samples were prepared the same way as described before. Briefly, 20 µl of 4x Laemmli buffer was added to ∼ 60 µl beads after removal of supernatant, followed by vortexing and then kept for shaking in a thermal mixer for 15 minutes at 95°C. Subsequently followed by another round of vortexing and shaking for additional 15 minutes at 95°C. After cooling the samples to room temperature, they were filtered using a 90 µm Mobi column filter. To reduce disulfide bridges, the 25 µl of the remaining were first diluted with 50 µl 50 mM HEPES solution and then treated with 2 µl 200 mM DTT in 50 mM HEPES at pH 8.5 for 30 min at 56°C. Carbamidomethylation of the accessible cysteine residues were done by adding 4 µl 400 mM 2-chloroacetamide in 50 mM HEPES at pH 8.5 and incubation for 30 min in the dark.

Protein clean up and digestion was done by SP3^44^. Sera-Mag Speed Beads (ThermoFisher) prewashed with water, 2 µl of a 1:1 mixture of the hydrophilic and hydrophobic beads was added to each sample at a concentration of 10 µg/µl. 83 µl acetonitrile was added to each sample and the suspensions were incubated for 8 min, subsequently the vials were kept on the magnetic stand for 2 more minutes. After discarding the supernatants, the beads were washed twice with 200 µl of 70% ethanol and once with 180 µl acetonitrile. After discarding the acetonitrile, the beads were air-dried. 150 µg trypsin in 10 µl of 50 mM HEPES buffer were added to the beads, the bead-bound proteins were digested for overnight. After incubation, on the next day, the bead suspensions were sonicated for 5 minutes and vortexed after which the vials were kept on the magnet. The supernatants containing the peptides were transferred to new vials. The magnetic beads were rinsed with 10 µl of 50 mM HEPES buffer and the resulting supernatants were combined with the first 10 µl. Following that individual samples were labelled by adding 4 µl TMT-6plex reagent (ThermoFisher) in acetonitrile, which were then incubated for one hour at room temperature. After one hour, the reactions were quenched with a solution of 5% hydroxylamine and then the mixture was acidified with 50 µl of 0.05% formic acid. The samples of obtained from each replicate were cleaned using an OASIS HLB µElution Plate (Waters). The wells were first washed twice with 0.05% formic acid in 80% acetonitrile and twice with 0.05% formic acid in water. The samples were loaded on the wells and washed with 0.05% formic acid in water. After elution with 0.05% formic acid in 80% acetonitrile the samples were dried and reconstituted in 4% acetonitrile and 1% formic acid in water.

#### Peptide fractionation

The pH of the samples was adjusted to 10 with ammonium hydroxide. The TMT-labelled peptides were then fractionated on an Agilent 1200 Infinity HPLC system equipped with a degasser, quaternary pump, autosampler, a variable wavelength UV detector (that was set to 254 nm), and fraction collector. Separation was performed on a Phenomenex Gemini C18 (100 x 1.0 mm; 3 μm; 110 Å) column using 20 mM ammonium formate at pH 10 in water as mobile phase A and 100% acetonitrile as mobile phase B. The column was used in combination with a Phenomenex Gemini C18, 4 x 2.0 mm Security Guard cartridge. The flow rate was 0.1 ml/min. After 2 min isocratic separation at 100% A, a linear gradient to 35% B at minute 59 was used, followed by washing at 85% B and reconstitution at 100% A. In total 32 two-minute fractions were collected and pooled. These were dried and reconstituted in 4% acetonitrile and 1% formic acid.

#### Mass spectrometry data acquisition

The fractionated samples were analyzed on an UltiMate 3000 nano LC system (Dionex) coupled to a QExactive plus (Thermo) mass spectrometer via a Nanospray Flex source (Thermo) using a Pico-Tip Emitter (New Objective; 360 µm OD x 20 µm ID; 10 µm tip). The peptides were first trapped on a C18 PepMap 100 µ-Precolumn (300 µm x 5 mm, 5 µm, 100 Å) prior to separation on a Waters nanoEase C18 75 µm x 250 mm, 1.8 µm, 100 Å column. The applied flow rates were 30 µl/min for trapping and 300 nl/min for separation. The mobile phase A was 0.1% formic acid in water and the mobile phase B was 0.1% formic acid in acetonitrile. After an initial isocratic step at 2% B for 2.9 minutes the multi-step gradient started with a gradient to 4% B at minute 4 followed by a linear increase to 8% B at minute 6. Subsequently, a shallow gradient to 28% B at minute 43 was followed by a steep gradient to 40% B at minute 52, a washing step at 80% B, and reconstitution at 2% B.

All spectra were acquired in positive ion mode. Full scan spectra were recorded in profile mode in a mass range of 375-1200 *m/z*, at a resolution of 70,000, with a maximum ion fill time of 10 ms and an AGC target value of 3x10^6^ ions. A top 20 method was applied with the normalized collision energy set to 32, an isolation window of 0.7, the resolution at 17,500, a maximum ion fill time of 50 ms, and an AGC target value of 2x10^5^ ions. The fragmentation spectra were recorded in profile mode with a fixed first mass of 100 *m/z*. Unassigned charge states as well as charge states of 1, 5-8, and >8 were excluded, and the dynamic exclusion was set to 30 seconds.

### Analysis of the MS results

#### MS data analysis for murine Th2 cells

IsobarQuant^45^ and Mascot (v2.2.07) were used to process the acquired data, which was searched against the Uniprot reference database of *Mus musculus* after the addition of common contaminants and reversed sequences. The following modifications were included into the search parameters: Carbamidomethyl (C) and TMT10 (K) (fixed modification), Acetyl (N-term), Oxidation (M) and TMT10 (N-term) (variable modifications). A mass error tolerance of 10 ppm was applied to full scan spectra and 0.02 Da to fragmentation spectra. Trypsin was selected as protease with an allowance of maximum two missed cleavages, a minimum peptide length of seven amino acids, and at least two unique peptides were required for protein identification. The false discovery rate (FDR) on peptide and protein level was set to 0.01.

The protein.txt output files from IsobarQuant were further processed using the R language. As quality filters, only proteins that were quantified with at least two unique peptides and have been identified in both biological replicates (analyzed in separate MS analysis) were used for further downstream analysis. The ‘signal_sum’ columns were used, and potential batch-effects were removed using the respective function from the limma package^46^. Subsequently, the data was normalized using a variance stabilization normalization^47^ (vsn). The limma package was employed again to test for differential abundance between the various experimental conditions. T-values of the limma output were pasted into fdrtool^48^ to estimate false discovery rates (q-values were used as FDR).

### Homology Modelling

The current study has been performed on five different protein targets. These are CYP51A1 (Uniprot Entry - Q8K0C4), LSS (Q8BLN5), GLUD1 (P26443), CLUH (Q5SW19), P4HB (P09103) and PITRM1 (Q8K411). The substrate selected is Pregnenolone (Pubchem compound CID: 8955). We have obtained our AlphaFold modelled structures for all the proteins from the UniProt database. AlphaFold^49, 50^ is a novel machine learning approach that incorporates physical and biological knowledge about protein structure, leveraging multi-sequence alignments, into the design of the deep learning algorithm. The state-of-art AI system developed by DeepMind, can computationally predict protein structures with unprecedented accuracy and speed. AlphaFold generated models have accuracy around 93% and can be used for purposeful drug designing related projects. AlphaFold identifiers selected for our studies are CYP51A1 (AF-Q8K0C4-F1), LSS (AF-Q8BLN5-F1), GLUD1 (AF-P26443-F1), CLUH (AF-Q5SW19-F1), P4HB (AF-P09103-F1) and PITRM1 (AF-Q8K411-F1).

### Molecular docking

AlphaFold protein models for CYP51A1, LSS, GLUD1, CLUH, P4HB and PITRM1 were prepared in MOE2020.0901 software (Molecular Operating Environment (MOE), 2020.0901; Chemical Computing Group ULC, 1010, Sherbooke St. West, Suite #910, Montreal, QC, Canada, H3A 2R7, 2022). The water molecules and heteroatoms were removed, and polar hydrogens were added. A temperature of 300K, salt concentration of 0.1 and pH 7 was quantified in an implicit solvated environment to undergo the protonation process. The structure was energy minimized in the Amber10: EHT force field to an RMS gradient of 0.01 kcal/mol/A^2. The energy minimized conformation of protein models was then subjected to 10 ns molecular dynamic simulations at a constant temperature of 300 K, heat time of 10 ps, and temperature relaxation of 0.2 ps to derive a stable conformation.

The ‘Site Finder’ feature on MOE 2020.0901 was used to predict the active sites of protein models. We have created a dummy model out of the generated active sites to derive the binding site of the modelled proteins. We have considered the largest active site based on participation of maximum amino acids for our docking studies. The docking process was conducted three times. The first and second docking experiment was performed by using ‘Rigid Receptor’ protocol. In this simulation, we considered ‘Triangle Matcher and London dG’^51^ as placement method and scores respectively. Similarly, “Rigid receptor/Induced Fit and GBVI-WSA dG’^51^ considered as refinement method and scores respectively. We have generated 30 poses for the placement method and 5 poses for the refinement method. The third docking process was carried out by using ‘Induced Fit’ protocol. In this step, the protein was made flexible to fit the conformation with the desired substrate. The rest of parameters under the current docking run remain the same as the previous docking simulation. At the end of the simulation, we chose the best substrate pose score according to their Gibbs free binding energy (ΔG binding), root mean square deviation (RMSD), and binding affinity between substrate and the protein models.

Depending on the settings used, the final result database contains the following fields: the final score (S); the root mean square deviation between the pose and the original substrate, or between the poses before and after refinement (rmsd and rmsd_refine, Å); energy of conformer (E_conf); and scores for successive docking stages: placement,rescoring, and refinement (E_place, E_score1, E_score2, E_refine). Lower final S-scores indicate more favourable poses. The detailed docking results of protein models are provided in Table: 1.

### Molecular dynamics and simulation

MD simulation was carried out to understand the stability and dynamic behaviour of modelled protein-substrate docked complexes. We used Desmond with Optimized Potentials for Liquid Simulations (OPLS4) as forcefield (Desmond, version 4.7., Schrodinger, LLC, New York, NY. 2022) in the experiments. The protein-substrate was saturated with SPC as solvent inside an orthorhombic box. It is further neutralized by adding appropriate counter ions and 0.15M of salt concentration^52^. The distance between the protein-substrate and box wall was set to 10Å to avoid the steric interaction. It was further minimised by applying a hybrid method of steepest descent and the Limited Memory Broyden–Fletcher–Goldfarb–Shanno (LBFGS) algorithms for 100ps until a gradient threshold of 25kcal/mol is achieved^53^. The temperature was maintained at 300K for whole simulations using Nose-Hoover thermostats. The Martyna-Tobias-Klein barostat method was used to maintain stable pressure. We performed NPT (Number of particles, Pressure and Temperature) ensemble for the equilibrated system for 100ns. To examine the equation of motion in dynamics, a multi-time step RESPA integrator algorithm was used^54^. The final equilibrated system is used to perform a 100ns molecular dynamics simulation run. The systems were relaxed by constant NVT (Number of particles, Volume and Temperature) ensemble conditions for 1ns to produce simulation data for post-simulation analysis^55, 56^ The results were analysed through simulation interaction diagram (SID) and trajectory plot module of Desmond.

### Binding Free Energy Calculation

The binding free energies of the protein-substrate complexes were calculated through Prime-MM/GBSA (Molecular Mechanics/Generalized Born Surface Area) (Prime, Schrodinger, LLC, New York, NY. 2022) to validate the Induced Fit docking results. MM/GBSA combines procedure that integrates OPLS molecular mechanics energies (EMM), an SGB solvation model for polar solvation (GSGB), and a non-polar solvation term (GNP). It comprises of nonpolar solvent accessible surface area and van der waals interactions^57^. The binding free energy calculation is expressed as:

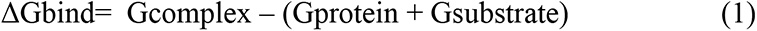

Where;

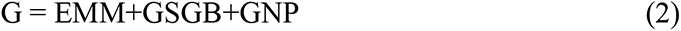

The Gaussian surface area model was preferred over vander waals by Prime (Prime, Schrodinger, LLC, New York, NY. 2022) for representing the solvent-accessible surface area^58^.

### Substrate Transport Analysis

Caver Web 1.0^59^ is used to study the comprehensive analysis of protein tunnels and channels for the substrate transport. The identified tunnels, their properties, energy profiles and trajectories for substrate passages can be calculated and visualized. The workflow involves 4 steps. The first step involves the selection of a protein structure and its pre-treatment. The second step is a selection of a starting point selection for tunnel detection. Protein tunnels are identified and analysed in the third step. The final step involves the selected substrate transport and the energy profile analysis for the selected tunnels.

## Author Contribution

SR designed the experiments. SR performed the experiments. SUR did the molecular docking and simulation studies. BM and JP provided the murine Th2 cells when required. SR wrote the manuscript with help from SUR. MLH did the on-bead digestion and mass spectrometry of CD4^+^ Th2 cells. SAT and A-CG supervised the study. All authors commented on and approved the draft manuscript before submission.

## Acknowledgments

SR acknowledges Ashoka University’s annual research funding for research visits. The ERC consolidator grant (ThDEFINE, Project ID: 646794) supported this study. A.-C.G. acknowledges the financial support of the Louis-Jeantet Foundation, Switzerland.

## Data and code availability

The TMT LC-MS/MS proteomics data have been deposited to the ProteomeXchange Consortium via the PRIDE^60^ partner repository with the dataset identifier PXD025574. Database: PXD025574.

