## Supplementary Figures for "CLICK- chemoproteomics and molecular dynamics simulation reveals pregnenolone targets and their binding conformations in Th2 cells"

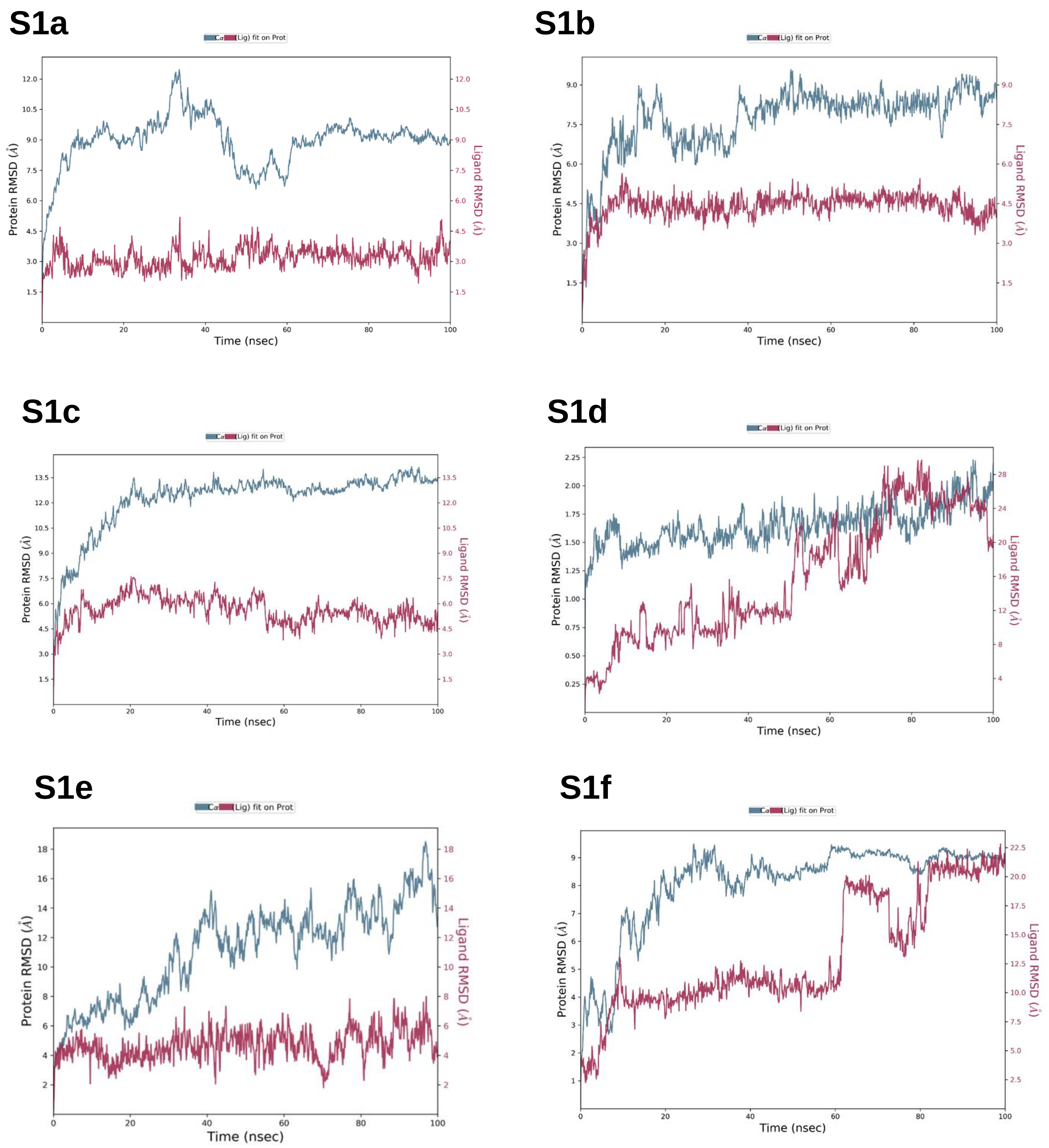


**Figure S1. Protein substrate RMSD** (a) P5-CLUH (b) P5-CYP51A1 (c) P5-GLUD1 (d) P5-LSS. (e) P5-P4HB. (f) P5-PITRM1

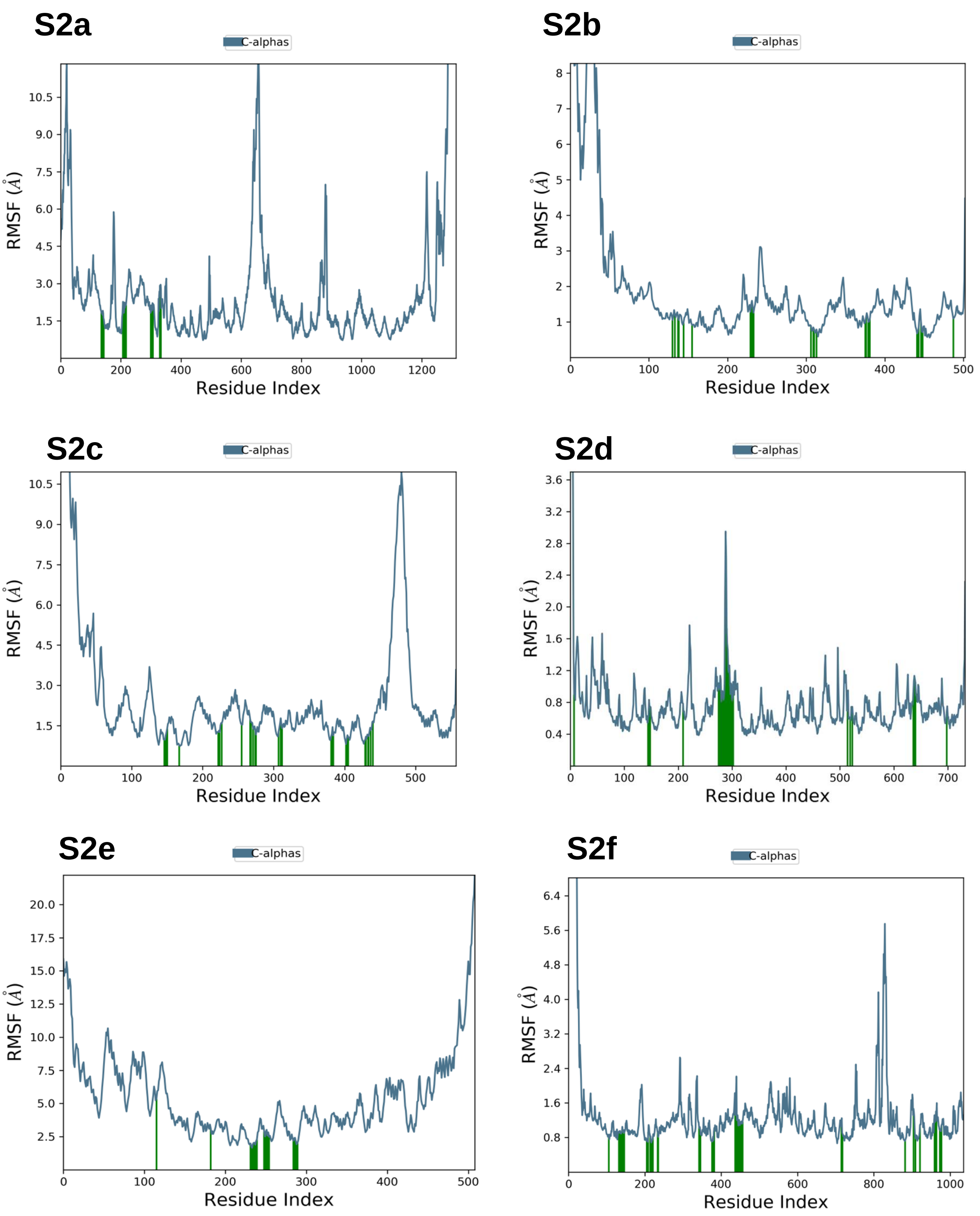


**Figure S2. Protein RMSF** (a) CLUH (b) CYP51A1 (c) GLUD1 (d) LSS. (e) P4HB. (f) PITRM1

S3a

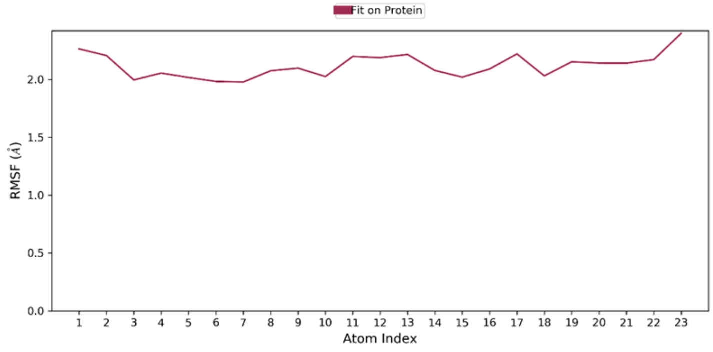


S3b

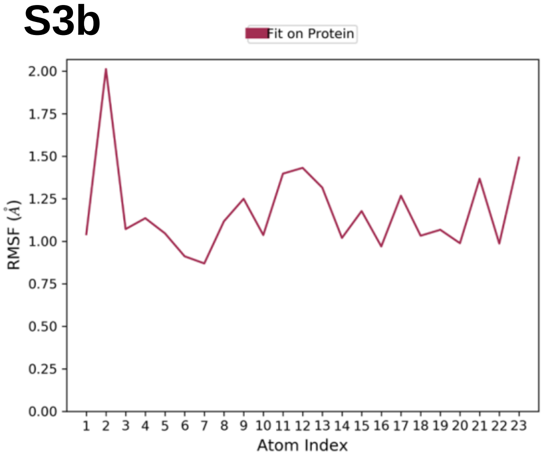


S3c

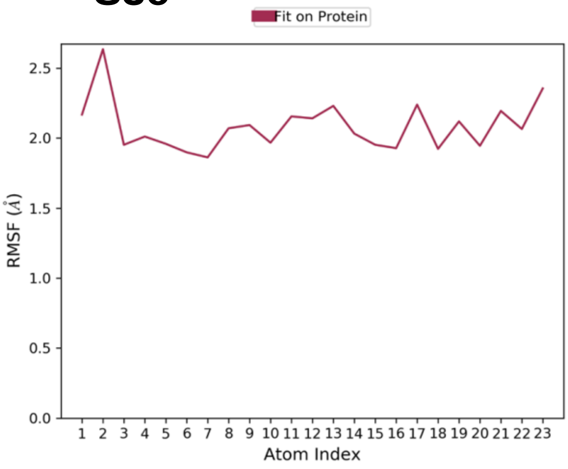


S3d

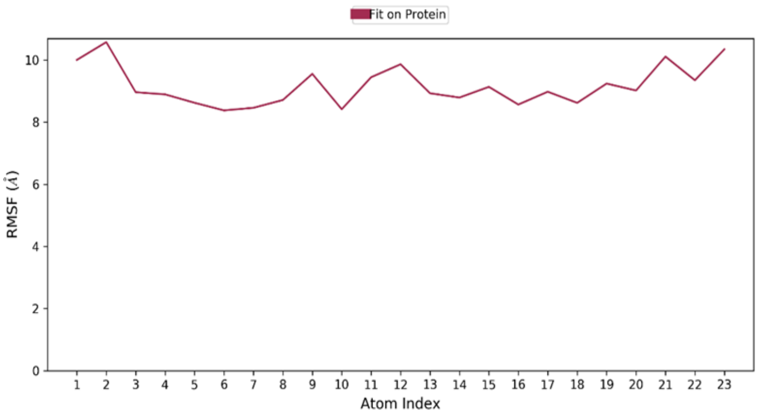


S3e

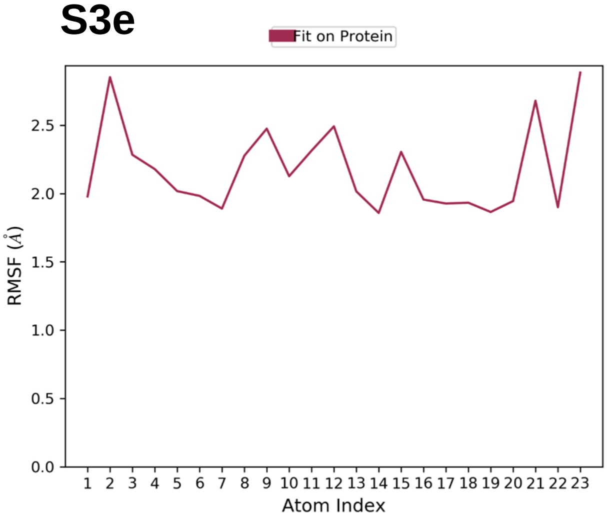


S3f

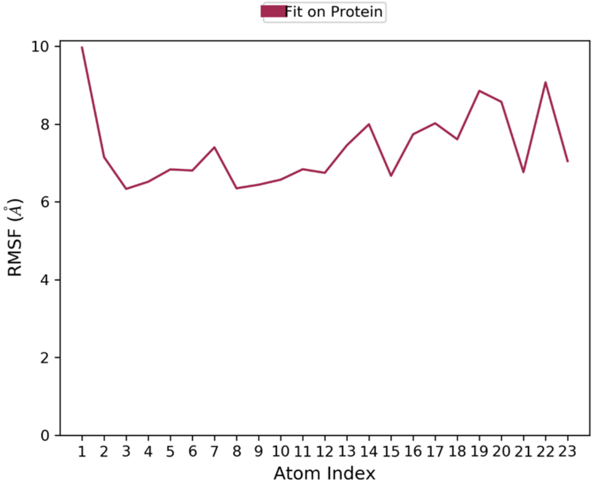


**Figure S3. Substrate (P5) RMSF** (a) CLUH (b) CYP51A1 (c) GLUD1 (d) LSS. (e) P4HB. (f) PITRM1

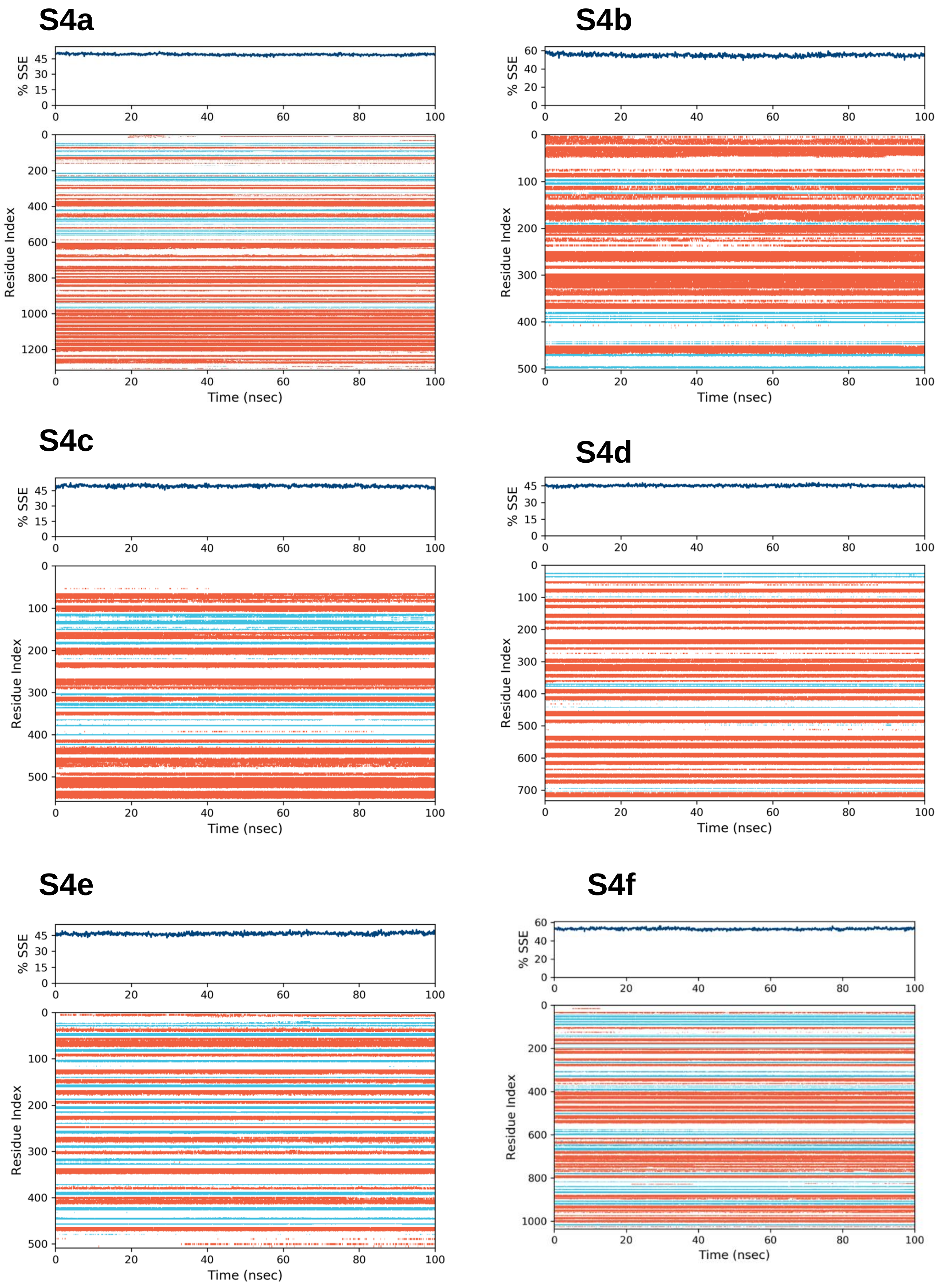


**Figure S4.** Secondary structure elements [SSE] (a) CLUH (b) CYP51A1 (c) GLUD1 (d) LSS. (e) P4HB. (f) PITRM1

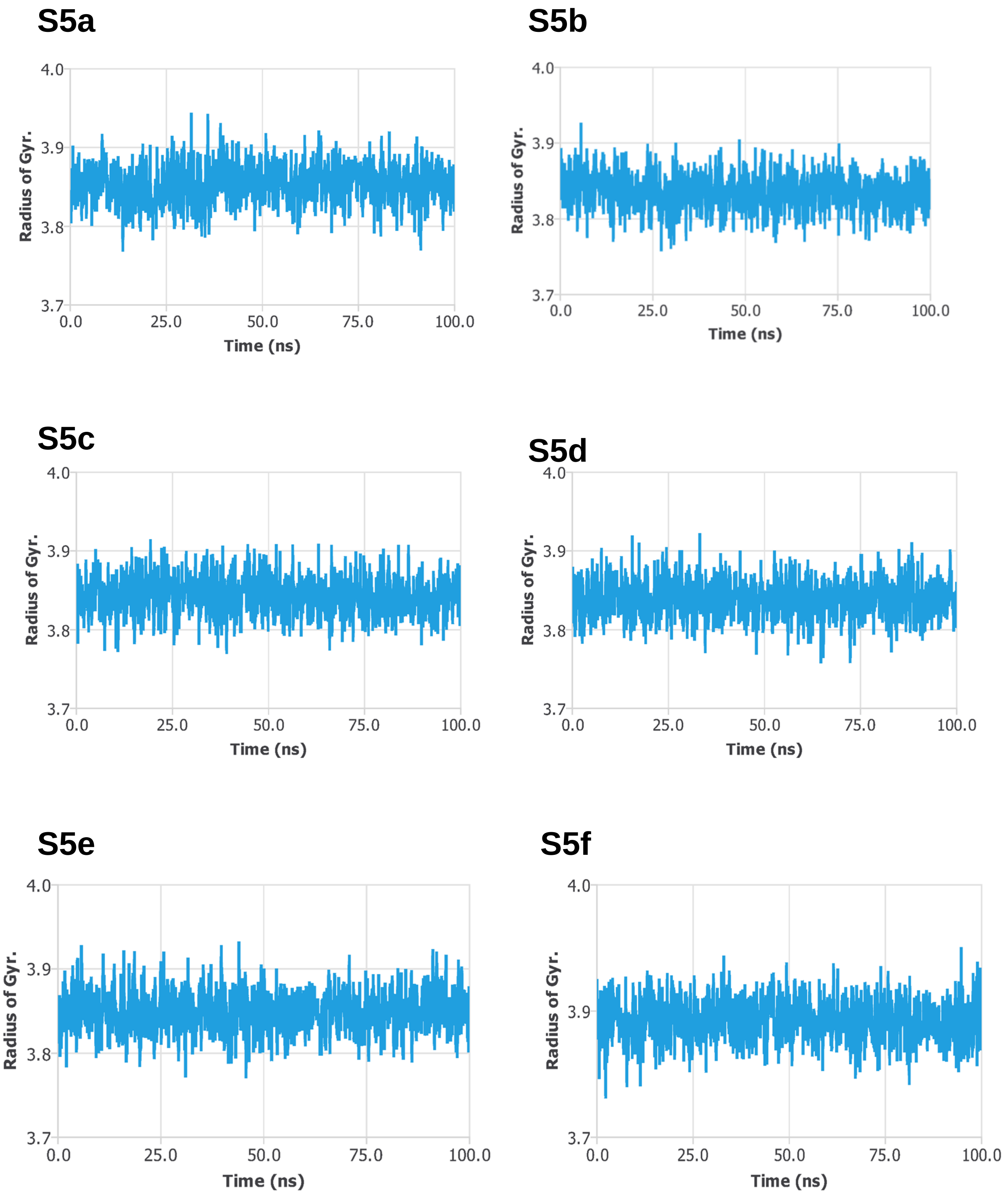


**Figure S5.** Radius of gyration (a) CLUH (b) CYP51A1 (c) GLUD1 (d) LSS. (e) P4HB. (f) PITRM1

**S6a**

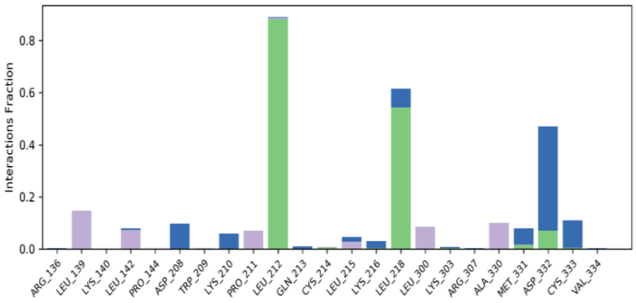


**S6b**

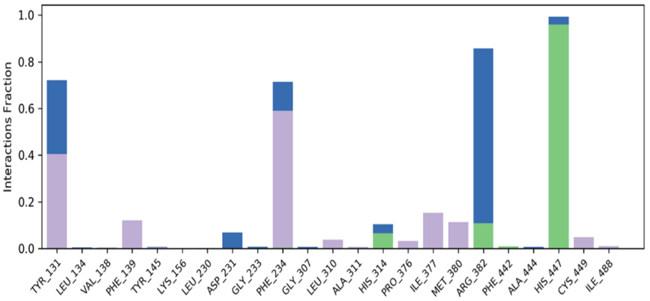


**S6c**

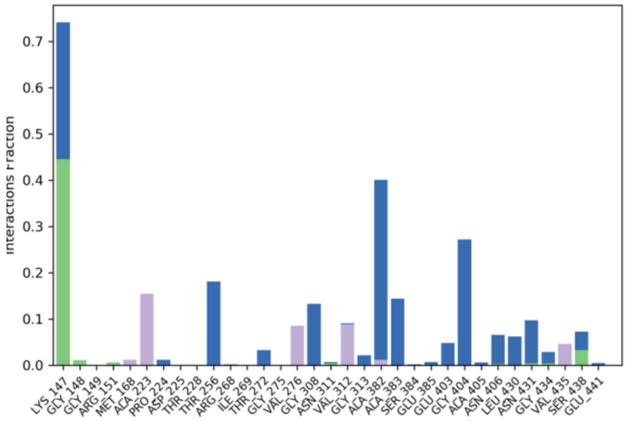


**S6d**

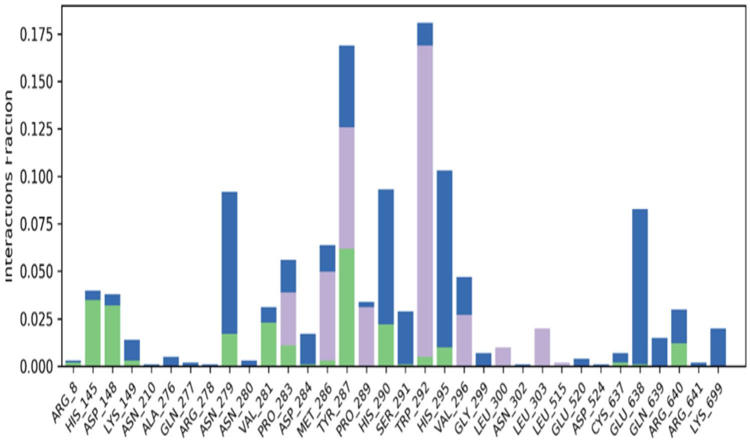


**S6e**

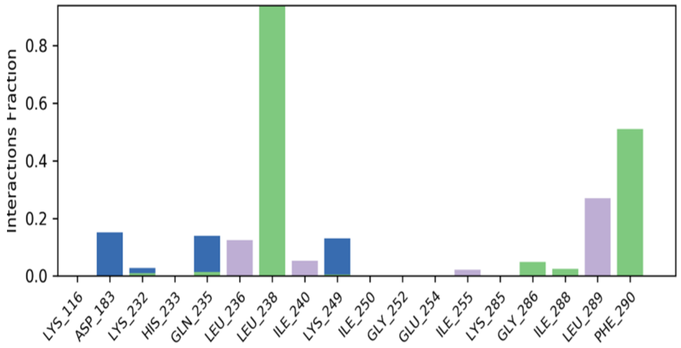


**S6f**

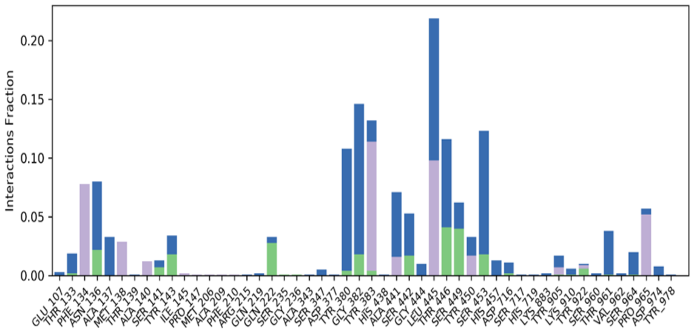


**Figure S6.** Protein - P5 contacts (a) CLUH (b) CYP51A1 (c) GLUD1 (d) LSS. (e) P4HB. (f) PITRM1

## S7a

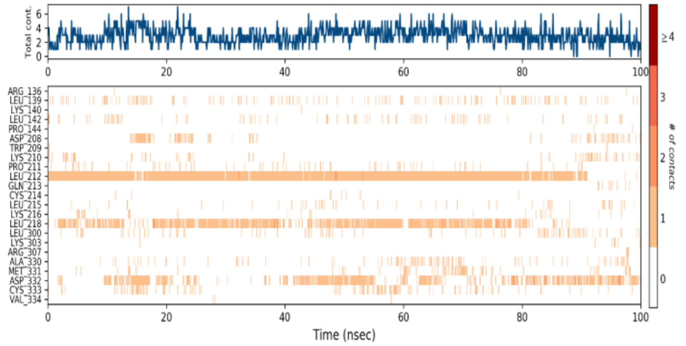


## S7b

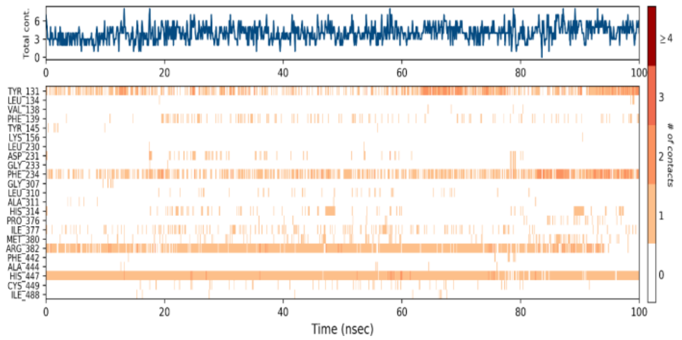


## S7c

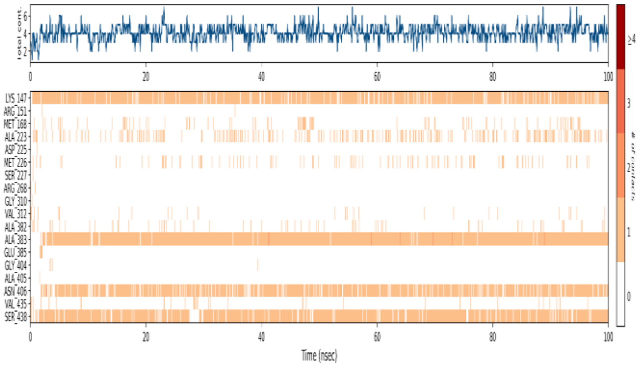


## S7d

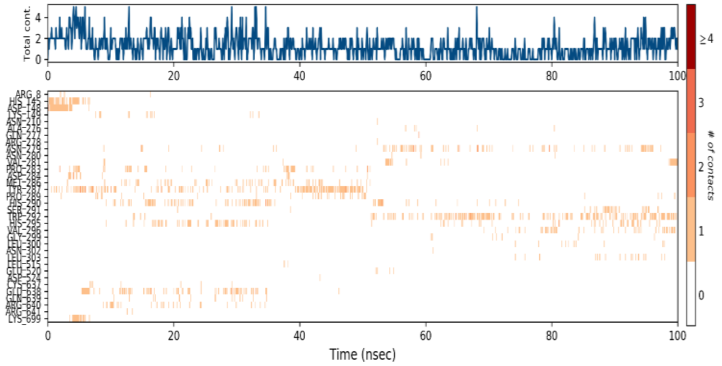


## S7e

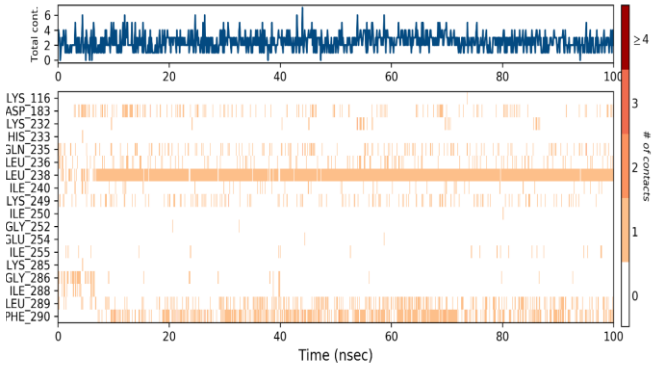


## S7f

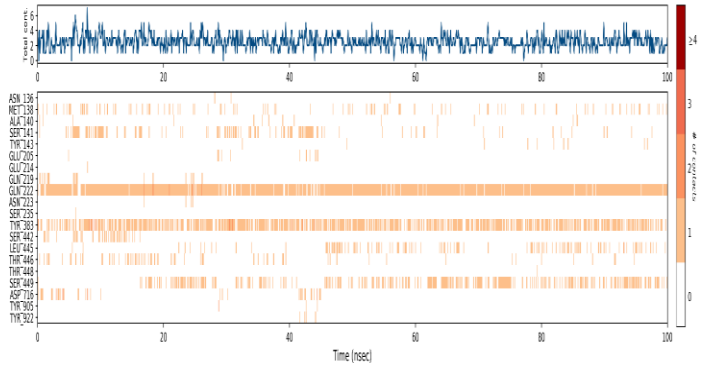


**Figure S7.** Protein - P5 timeline (a) CLUH (b) CYP51A1 (c) GLUD1 (d) LSS. (e) P4HB. (f) PITRM1

**S8a**

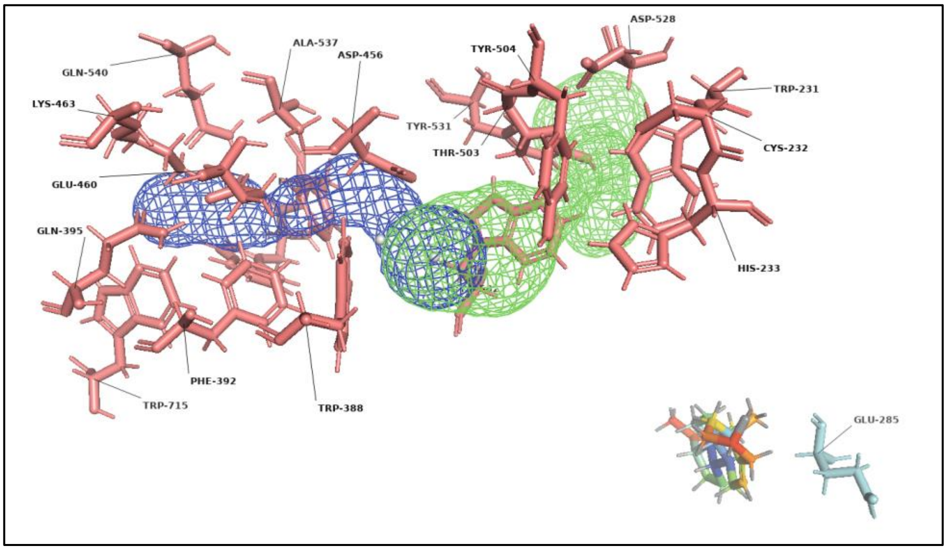


**S8b**

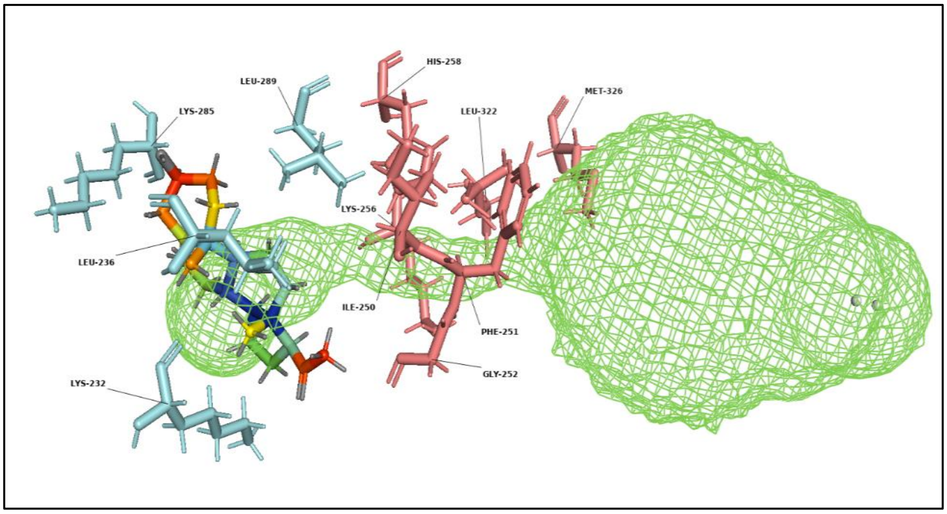


**S8c**

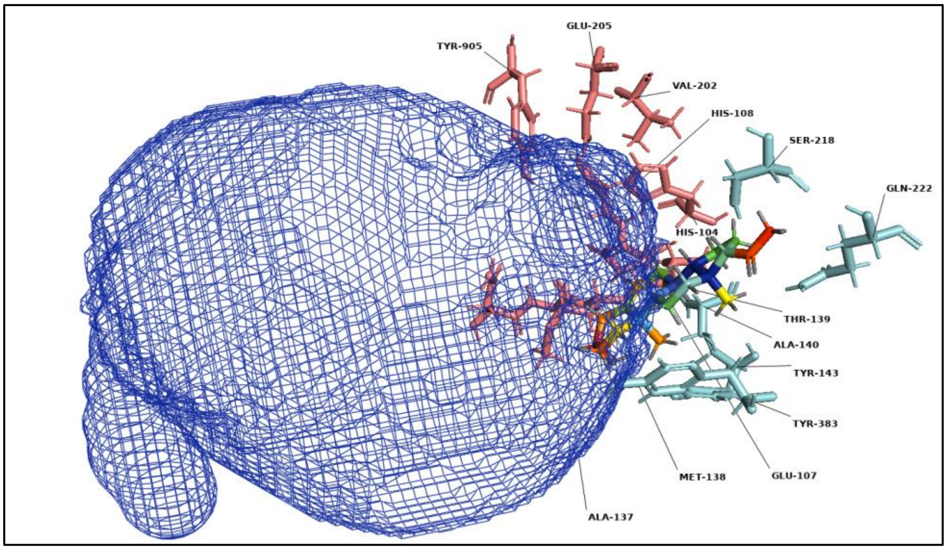


**Figure S8.** Non-involvement of tunnels for P5 transport. (a) LSS protein, Cyan color denotes hydrogen and hydrophobic interactions; Red Salmon color denotes other amino acids surrounding the tunnel. (b) P4HB protein, Cyan color denotes hydrogen and hydrophobic interactions; Red Salmon color denotes other amino acids surrounding the tunnel. (C) PITRM1 protein, Cyan color denotes hydrogen and hydrophobic interactions; Red Salmon color denotes other amino acids surrounding the tunnel. (b)
