## Supplementary Tables for "CLICK- chemoproteomics and molecular dynamics simulation reveals pregnenolone targets and their binding conformations in Th2 cells"

**Protein-Ligand Interaction Profiler**

**Table ST1(a):** PLIF results of CLUH

| Proteins | **Hydrophobic interactions** |  |  |  |  | |  | | |  |  |  |
| --- | --- | --- | --- | --- | --- | --- | --- | --- | --- | --- | --- | --- |
|  | Index | Residue | AA | Distance | Substrate Atom | | Protein Atom | | |  |  |  |
| **CLUH** | 1 | 143A | ASP | 3.91 | 20723 | | 2213 | | |  |  |  |
|  | 2 | 332A | ASP | 3.80 | 20728 | | 5204 | | |  |  |  |
| Proteins | **Hydrogen Bonds** |  |  |  |  |  | |  |  | |  |  |
|  | Index | Residues | AA | Distance H-A | Distance D-A | Donor Angle | | Protein Donor | Side Chain | | Donor Atom | Acceptor Atom |
| **CLUH** | 1 | 212A | LEU | 2.42 | 2.96 | 113.15 | | YES | NO | | 3253[Nam] | 20713[O2] |
|  | 2 | 218A | LEU | 2.17 | 2.99 | 137.32 | | YES | NO | | 3357[Nam] | 20712[O3] |

**Table ST1(b):** PLIF results of CYP51A1

| Proteins | | **Hydrophobic interactions** | |  | |  | |  | |  | | |  | |  |  |  |
| --- | --- | --- | --- | --- | --- | --- | --- | --- | --- | --- | --- | --- | --- | --- | --- | --- | --- |
|  | | Index | | Residue | | AA | | Distance | | Substrate Atom | | | Protein Atom | |  |  |  |
| **CYP51A1** | | 1 | | 131A | | TYR | | 3.70 | | 8055 | | | 1989 | |  |  |  |
|  | | 2 | | 131A | | TYR | | 3.65 | | 8054 | | | 1991 | |  |  |  |
|  | | 3 | | 131A | | TYR | | 3.67 | | 8056 | | | 1992 | |  |  |  |
|  | | 4 | | 145A | | TYR | | 3.59 | | 8055 | | | 2203 | |  |  |  |
|  | | 5 | | 234A | | PHE | | 3.49 | | 8052 | | | 3629 | |  |  |  |
|  | | 6 | | 234A | | PHE | | 3.54 | | 8060 | | | 3630 | |  |  |  |
|  | | 7 | | 377A | | ILE | | 3.72 | | 8054 | | | 5982 | |  |  |  |
|  | | 8 | | 377A | | ILE | | 3.92 | | 8043 | | | 5983 | |  |  |  |
|  | | 9 | | 487A | | MET | | 3.86 | | 8049 | | | 7732 | |  |  |  |
|  | | 10 | | 488A | | ILE | | 3.78 | | 8049 | | | 7753 | |  |  |  |
| Proteins | **Hydrogen Bonds** | |  | |  | |  | |  | |  |  | |  | |  |  |
|  | Index | | Residue | | AA | | Distance H-A | | Distance D-A | | Donor Angle | Protein Donor | | Side Chain | | Donor Atom | Acceptor Atom |
| **CYP51A1** | 1 | | 145A | | TYR | | 3.28 | | 3.64 | | 104.61 | YES | | YES | | 2204[O3] | 8038[O3] |
|  | 2 | | 145A | | TYR | | 2.73 | | 3.64 | | 160.21 | NO | | YES | | 8038[O3] | 2204[O3] |
|  | 3 | | 231A | | ASP | | 3.21 | | 3.51 | | 101.27 | YES | | YES | | 3601[O3] | 8039[O2] |

**Table ST1(c):** PLIF results of GLUD1

| Proteins | | **Hydrophobic interactions** | |  | |  | |  | |  | | |  | |  |  |  |
| --- | --- | --- | --- | --- | --- | --- | --- | --- | --- | --- | --- | --- | --- | --- | --- | --- | --- |
|  | | Index | | Residue | | AA | | Distance | | Substrate Atom | | | Protein Atom | |  |  |  |
| **GLUD1** | | 1 | | 311A | | ASN | | 3.87 | | 8614 | | | 4726 | |  |  |  |
| Proteins | **Hydrogen Bonds** | |  | |  | |  | |  | |  |  | |  | |  |  |
|  | Index | | Residue | | AA | | Distance H-A | | Distance D-A | | Donor Angle | Protein Donor | | Side Chain | | Donor Atom | Acceptor Atom |
| **GLUD1** | 1 | | 268A | | ARG | | 3.06 | | 3.79 | | 130.07 | YES | | YES | | 4094[Ng+] | 8605[O3] |
|  | 2 | | 268A | | ARG | | 2.23 | | 3.17 | | 153.56 | YES | | YES | | 4095[Ng+] | 8605[O3] |
|  | 3 | | 272A | | THR | | 2.88 | | 3.41 | | 115.63 | YES | | YES | | 4156[O3] | 8605[O3] |
|  | 4 | | 272A | | THR | | 2.93 | | 3.41 | | 112.11 | NO | | YES | | 8605[O3] | 4156[O3] |

**Table ST1(d):** PLIF results of LSS

| Proteins | | **Hydrophobic interactions** | |  | |  | |  | |  | | |  | |  |  |  |
| --- | --- | --- | --- | --- | --- | --- | --- | --- | --- | --- | --- | --- | --- | --- | --- | --- | --- |
|  | | Index | | Residue | | AA | | Distance | | Substrate Atom | | | Protein Atom | |  |  |  |
| **LSS** | | 1 | | 285A | | GLU | | 3.96 | | 11505 | | | 4472 | |  |  |  |
| Proteins | **Hydrogen Bonds** | |  | |  | |  | |  | |  |  | |  | |  |  |
|  | Index | | Residue | | AA | | Distance H-A | | Distance D-A | | Donor Angle | Protein Donor | | Side Chain | | Donor Atom | Acceptor Atom |
| **LSS** | 1 | | 285A | | GLU | | 3.29 | | 3.78 | | 114.36 | NO | | NO | | 11487[O3] | 4473[O2] |

**Table ST1(e):** PLIF results of P4HB

| Proteins | **Hydrophobic interactions** |  |  |  |  |  |
| --- | --- | --- | --- | --- | --- | --- |
|  | Index | Residue | AA | Distance | Substrate Atom | Protein Atom |
| **P4H8** | 1 | 232A | LYS | 3.57 | 7984 | 3557 |
|  | 2 | 236A | LEU | 3.37 | 7980 | 3628 |
|  | 3 | 285A | LYS | 3.36 | 7991 | 4409 |
|  | 4 | 289A | LEU | 3.49 | 7992 | 4480 |

**Table ST1(f):** PLIF results of PITRM1

| Proteins | | **Hydrophobic interactions** | |  | |  | |  | |  | | |  | |  |  |  |
| --- | --- | --- | --- | --- | --- | --- | --- | --- | --- | --- | --- | --- | --- | --- | --- | --- | --- |
|  | | Index | | Residue | | AA | | Distance | | Substrate Atom | | | Protein Atom | |  |  |  |
| **PITRM1** | | 1 | | 140A | | ALA | | 3.86 | | 16455 | | | 2246 | |  |  |  |
|  | | 2 | | 222A | | GLN | | 3.64 | | 16460 | | | 3584 | |  |  |  |
|  | | 3 | | 383A | | TYR | | 3.79 | | 16452 | | | 6069 | |  |  |  |
|  | | 4 | | 383A | | TYR | | 3.65 | | 16447 | | | 6073 | |  |  |  |
|  | | 5 | | 383A | | TYR | | 3.84 | | 16453 | | | 6074 | |  |  |  |
|  | | 6 | | 383A | | TYR | | 3.50 | | 16455 | | | 6072 | |  |  |  |
| Proteins | **Hydrogen Bonds** | |  | |  | |  | |  | |  |  | |  | |  |  |
|  | Index | | Residue | | AA | | Distance H-A | | Distance D-A | | Donor Angle | Protein Donor | | Side Chain | | Donor Atom | Acceptor Atom |
| **PITRM1** | 1 | | 143A | | TYR | | 2.90 | | 3.73 | | 145.47 | YES | | YES | | 2286[O3] | 16438[O3] |
|  | 2 | | 143A | | TYR | | 3.15 | | 3.73 | | 120.23 | NO | | YES | | 16438[O3] | 2286[O3] |
|  | 3 | | 218A | | SER | | 3.70 | | 4.03 | | 102.98 | YES | | YES | | 3522[O3] | 16439[O2] |
|  | 4 | | 222A | | GLN | | 2.68 | | 3.12 | | 106.49 | YES | | YES | | 3588[Nam] | 16439[O2] |

**Table ST2:** MM-GBSA residue decomposition results for the selected proteins

| Proteins |  |  |  |  |  |  |  |  |  |
| --- | --- | --- | --- | --- | --- | --- | --- | --- | --- |
| 100 ns | Dg bind | Coulomb | Solvation | Covalent | Vdw | H-bond | Lipo | Pi-Pi | Contact |
| **CLUH** | **-30.31** | **-6.2** | **7.92** | **0** | **-24.04** | **-0.44**  **LEU212**  **[-0.24], LYS216**  **[-0.01], LEU218**  **[-0.14], LYS303**  **[-0.01], ASP332**  **[-0.03]** | **-8.43** | **0** | **0** |
| GLUD1 | -16.35 | -2.31 | 8.9 | 0 | -18.58 | -0.32  LYS147  [-0.15], GLY148  [-0.01], THR256  [-0.03], ARG268  [-0.03], GLY434  [-0.01], SER438  [-0.08] | -5.64 | 0 | 0 |
| **CYP51A1** | **-30.69** | **-4.42** | **9.96** | **0** | **-24.58** | **-0.38 HIS314**  **[-0.03], ARG382**  **[-0.08], HIS447**  **[-0.28]** | **-13.02** | **0** | **0** |
| LSS | -15.48 | -2.16 | 3.93 | 0 | -12.46 | -0.11 ASP148  [-0.03], ASN279  [-0.01], TYR287  [-0.05], ARG640  [-0.02] | -5.77 | 0 | 0 |
| **P4H8** | **-30.02** | **-8.68** | **8.89** | **0** | **-23.36** | **-0.54 (LYS232**  **[-0.01], GLN235**  **[-0.01], LEU238**  **[-0.24], GLY286**  **[-0.02], ILE288**  **[-0.01], PHE290**  **[-0.24]** | **-8.85** | **0** | **0** |
| PITRM1 | -14.23 | -2.44 | 6.56 | 0 | -12.91 | -0.11 ASN136  [-0.02], TYR143  [-0.01] TYR380  [-0.01], GLY382  [-0.02], TYR383  [-0.01], THR446  [-0.01], SER453  [-0.03] | -6.36 | 0 | 0 |

**Table ST3:** Tunnels and channels detected in CLUH, CYP51A1, LSS, GLUD1, P4HB and PITRM1

**Tunnel**-identifier of protein tunnel; **Bottleneck radius**-radius of the narrowest part of the tunnel; **Length**-Length of the tunnel; **Curvature**-curvature of the tunnel; **Throughput**-throughput of the tunnel

| **Details of Tunnel** | **S. No** |  |  |  |  |  |  |  |  |  |
| --- | --- | --- | --- | --- | --- | --- | --- | --- | --- | --- |
| **GLUD1** | **Tunnel** | **Bottleneck Radius [A˚]:** | **Length**  **[A˚]:** | **Distance to surface**  **[A˚]:** | **Distance from starting point [A˚]:** | **Curvature** | **Throughput** | **Number of Residues** | **Number of Bottleneck** | **Bottleneck Residues (12)** |
|  | 2 | 1.8 | 9.2 | 7.9 | 0.5 | 1.2 | 0.77 | 23 | 1 | **ALA223, PRO224, ASP225, GLY229,**  **GLY257,**  **ARG268, ILE269, THR272, ASN311, VAL312, GLY434, THR228** |
| **CYP51A1** | **Tunnel** | **Bottleneck Radius [A˚]:** | **Length**  **[A˚]:** | **Distance to surface**  **[A˚]:** | **Distance from starting point [A˚]:** | **Curvature** | **Throughput** | **Number of Residues** | **Number of Bottleneck** | **Bottleneck Residues (7)** |
|  | 1 | 1.7 | 9.7 | 8.4 | 7.5 | 1.2 | 0.79 | 22 | 1 | **ASP231, PHE234, THR235, HIS314, THR486, MET487, ILE488** |
|  |  |  |  |  |  |  |  |  |  | **Bottleneck Residues (9)** |
|  | 2 | 2.0 | 15.9 | 12.7 | 4.5 | 1.3 | 0.76 | 33 | 1 | **TYR131, LEU134, ILE377, MET378, THR379, MET380, MET381, MET487, ILE488** |
|  | **Tunnel** | **Bottleneck Radius [A˚]:** | **Length**  **[A˚]:** | **Distance to surface**  **[A˚]:** | **Distance from staring point [A˚]:** | **Curvature** | **Throughput** | **Number of Residues** | **Number of Bottleneck** | **Bottleneck Residues (7)** |
| **CLUH** | 6 | 1.0 | 24.1 | 12.4 | 8.5 | 1.9 | 0.45 | 34 | 1 | **LYS140, ASP143, PRO144, SER145, ASP146, ALA147, ASN154** |
| **PITRM1**  **(No bottleneck amino acid reported for substrate binding)** | **Tunnel** | **Bottleneck Radius [A˚]:** | **Length**  **[A˚]:** | **Distance to surface**  **[A˚]:** | **Distance from staring point [A˚]:** | **Curvature** | **Throughput** | **Number of Residues** | **Number of Bottleneck** | **Bottleneck Residues (10)** |
|  | 1 | 1.6 | 51.6 | 28.4 | 0.0 | 1.8 | 0.67 | 76 | 1 | **HIS104, GLU107, HIS108, ASN136, ALA137, MET138, THR139, VAL202, GLU205, TYR905** |
| **LSS**  **(No bottleneck amino acid reported for substrate binding)** | **Tunnel** | **Bottleneck Radius [A˚]:** | **Length**  **[A˚]:** | **Distance to surface**  **[A˚]:** | **Distance from staring point [A˚]:** | **Curvature** | **Throughput** | **Number of Residues** | **Number of Bottleneck** | **Bottleneck Residues (11)** |
|  | 1 | 1.0 | 17.1 | 13.9 | 11.0 | 1.2 | 0.44 | 33 | 1 | **TRP388, PHE392, GLN395, ASP456, GLU460, LYS463, ALA537, GLN540, TRP591, PHE592, TRP715** |
|  | **Tunnel** | **Bottleneck Radius [A˚]:** | **Length**  **[A˚]:** | **Distance to surface**  **[A˚]:** | **Distance from staring point [A˚]:** | **Curvature** | **Throughput** | **Number of Residues** | **Number of Bottleneck** | **Bottleneck Residues (8)** |
|  | 2 | 0.9 | 26.2 | 15.6 | 8.0 | 1.7 | 0.33 | 36 | 1 | **TRP231, CYS232, HIS233, THR503, TYR504, ASP528, TYR531, TRP582** |
| **P4H8**  **(No bottleneck amino acid reported for substrate binding)** | **Tunnel** | **Bottleneck Radius [A˚]:** | **Length**  **[A˚]:** | **Distance to surface**  **[A˚]:** | **Distance from staring point [A˚]:** | **Curvature** | **Throughput** | **Number of Residues** | **Number of Bottleneck** | **Bottleneck Residues (7)** |
|  | 22 | 1.0 | 42.8 | 30.9 | 23.0 | 1.4 | 0.30 | 42 | 1 | **ILE250, PHE251, GLY252, LYS256, HIS258, LEU322, MET326** |

**Table ST4:** Atoms positioning of amino acids inside the tunnel during bond formation with the substrate for GLUD1, CLUH, CYP51A1

| **CYP51A1** | **S No:** | **ID** | **Name** |
| --- | --- | --- | --- |
|  | Tunnel |  |  |
| **ASP231** | 1 | 3596 | CA |
|  |  | 3597 | C |
|  |  | 3598 | CB |
|  |  | 3599 | O |
|  |  | 3600 | CG |
|  |  | 3601 | OD1 |
|  |  | 3602 | OD2 |
|  |  | 3604 | HA |
|  |  | 3605 | HB2 |
|  |  | 3606 | HB3 |
| **PHE234** | 1 | 3621 | N |
|  |  | 3622 | CA |
|  |  | 3623 | C |
|  |  | 3624 | CB |
|  |  | 3625 | O |
|  |  | 3626 | CG |
|  |  | 3627 | CD1 |
|  |  | 3628 | CD2 |
|  |  | 3629 | CE1 |
|  |  | 3630 | CE2 |
|  |  | 3631 | CZ |
|  |  | 3632 | H |
|  |  | 3633 | HA |
|  |  | 3634 | HB2 |
|  |  | 3635 | HB3 |
|  |  | 3636 | HD1 |
|  |  | 3637 | HD2 |
|  |  | 3638 | HE1 |
|  |  | 3639 | HE2 |
|  |  | 3640 | HZ |
| **MET487** | 1 | 7729 | N |
|  |  | 7730 | CA |
|  |  | 7731 | C |
|  |  | 7732 | CB |
|  |  | 7733 | O |
|  |  | 7734 | CG |
|  |  | 7735 | SD |
|  |  | 7736 | CE |
|  |  | 7737 | H |
|  |  | 7738 | HA |
|  |  | 7739 | HB2 |
|  |  | 7740 | HB3 |
|  |  | 7741 | HG2 |
|  |  | 7742 | HG3 |
|  |  | 7743 | HE1 |
|  |  | 7744 | HE2 |
|  |  | 7745 | HE3 |
| **ILE488** | 1 | 7746 | N |
|  |  | 7747 | CA |
|  |  | 7748 | C |
|  |  | 7749 | CB |
|  |  | 7750 | O |
|  |  | 7751 | CG1 |
|  |  | 7752 | CG2 |
|  |  | 7753 | CD1 |
|  |  | 7754 | H |
|  |  | 7755 | HA |
|  |  | 7756 | HB |
|  |  | 7757 | HG12 |
|  |  | 7758 | HG13 |
|  |  | 7761 | HD23 |
|  |  | 7762 | HD11 |
|  |  | 7763 | HD12 |
|  |  | 7764 | HD13 |
| **TYR131** | 2 | 1984 | CA |
|  |  | 1985 | C |
|  |  | 1986 | CB |
|  |  | 1987 | O |
|  |  | 1988 | CG |
|  |  | 1989 | CD1 |
|  |  | 1990 | CD2 |
|  |  | 1991 | CE1 |
|  |  | 1992 | CE2 |
|  |  | 1993 | OH |
|  |  | 1994 | CZ |
|  |  | 1996 | HA |
|  |  | 1997 | HB2 |
|  |  | 1998 | HB3 |
|  |  | 1999 | HD1 |
|  |  | 2000 | HD2 |
|  |  | 2001 | HE1 |
|  |  | 2002 | HE2 |
|  |  | 2003 | HH |
| **ILE377** | 2 | 5976 | N |
|  |  | 5977 | CA |
|  |  | 5978 | C |
|  |  | 5979 | CB |
|  |  | 5980 | O |
|  |  | 5981 | CG1 |
|  |  | 5982 | CG2 |
|  |  | 5983 | CD1 |
|  |  | 5985 | HA |
|  |  | 5986 | HB |
|  |  | 5987 | HG12 |
|  |  | 5988 | HG13 |
|  |  | 5989 | HG21 |
|  |  | 5990 | HG22 |
|  |  | 5991 | HG23 |
|  |  | 5992 | HD11 |
|  |  | 5993 | HD12 |
|  |  | 5994 | HD13 |
| **MET487** | 2 | 7729 | N |
|  |  | 7730 | CA |
|  |  | 7731 | C |
|  |  | 7732 | CB |
|  |  | 7733 | O |
|  |  | 7734 | CG |
|  |  | 7735 | SD |
|  |  | 7736 | CE |
|  |  | 7737 | H |
|  |  | 7738 | HA |
|  |  | 7739 | HB2 |
|  |  | 7740 | HB3 |
|  |  | 7741 | HG2 |
|  |  | 7742 | HG3 |
|  |  | 7743 | HE1 |
|  |  | 7744 | HE2 |
|  |  | 7745 | HE3 |
| **ILE488** | 2 | 7746 | N |
|  |  | 7747 | CA |
|  |  | 7748 | C |
|  |  | 7749 | CB |
|  |  | 7750 | O |
|  |  | 7751 | CG1 |
|  |  | 7752 | CG2 |
|  |  | 7753 | CD1 |
|  |  | 7754 | H |
|  |  | 7755 | HA |
|  |  | 7756 | HB |
|  |  | 7757 | HG12 |
|  |  | 7758 | HG13 |
|  |  | 7761 | HD23 |
|  |  | 7762 | HD11 |
|  |  | 7763 | HD12 |
|  |  | 7764 | HD13 |
| **GLUD1** |  |  |  |
| **ARG268** | 2 | 4087 | CA |
|  |  | 4088 | C |
|  |  | 4089 | CB |
|  |  | 4090 | O |
|  |  | 4091 | CG |
|  |  | 4092 | CD |
|  |  | 4093 | NE |
|  |  | 4094 | NH1 |
|  |  | 4095 | NH2 |
|  |  | 4096 | CZ |
|  |  | 4101 | HG2 |
|  |  | 4102 | HG3 |
|  |  | 4104 | HD3 |
|  |  | 4105 | HE |
|  |  | 4106 | HH11 |
|  |  | 4107 | HH12 |
|  |  | 4108 | HH21 |
|  |  | 4109 | HH22 |
| **THR272** | 2 | 4150 | N |
|  |  | 4151 | CA |
|  |  | 4153 | CB |
|  |  | 4155 | CG2 |
|  |  | 4156 | OG1 |
|  |  | 4157 | H |
|  |  | 4159 | HB |
|  |  | 4160 | HG21 |
|  |  | 4161 | HG22 |
|  |  | 4162 | HG23 |
|  |  | 4163 | HG1 |
| **ASN311** | 2 | 4723 | N |
|  |  | 4724 | CA |
|  |  | 4725 | C |
|  |  | 4726 | CB |
|  |  | 4727 | O |
|  |  | 4728 | CG |
|  |  | 4729 | ND2 |
|  |  | 4730 | OD1 |
|  |  | 4731 | H |
|  |  | 4732 | HA |
|  |  | 4733 | HB2 |
|  |  | 4734 | HB3 |
|  |  | 4735 | HD21 |
|  |  | 4736 | HD22 |
| **CLUH** |  |  |  |
| **ASP143** | 6 | 2210 | N |
|  |  | 2211 | CA |
|  |  | 2212 | C |
|  |  | 2213 | CB |
|  |  | 2214 | O |
|  |  | 2215 | CG |
|  |  | 2216 | OD1 |
|  |  | 2217 | OD2 |
|  |  | 2218 | H |
|  |  | 2219 | HA |
|  |  | 2220 | HB2 |
|  |  | 2221 | HB3 |
